# A recurrent neural circuit in *Drosophila* deblurs visual inputs

**DOI:** 10.1101/2024.04.19.590352

**Authors:** Michelle M. Pang, Feng Chen, Marjorie Xie, Shaul Druckmann, Thomas R. Clandinin, Helen H. Yang

## Abstract

A critical goal of vision is to detect changes in light intensity, even when these changes are blurred by the spatial resolution of the eye and the motion of the animal. Here we describe a recurrent neural circuit in *Drosophila* that compensates for blur and thereby selectively enhances the perceived contrast of moving edges. Using *in vivo*, two-photon voltage imaging, we measured the temporal response properties of L1 and L2, two cell types that receive direct synaptic input from photoreceptors. These neurons have biphasic responses to brief flashes of light, a hallmark of cells that encode changes in stimulus intensity. However, the second phase was often much larger than the first, creating an unusual temporal filter. Genetic dissection revealed that recurrent neural circuitry strongly shapes the second phase of the response, informing the structure of a dynamical model. By applying this model to moving natural images, we demonstrate that rather than veridically representing stimulus changes, this temporal processing strategy systematically enhances them, amplifying and sharpening responses. Comparing the measured responses of L2 to model predictions across both artificial and natural stimuli revealed that L2 tunes its properties as the model predicts in order to deblur images. Since this strategy is tunable to behavioral context, generalizable to any time-varying sensory input, and implementable with a common circuit motif, we propose that it could be broadly used to selectively enhance sharp and salient changes.

## Introduction

Determining how the intensity of light changes over space and time is fundamental to vision. However, both the spatial acuity of the eye and movements of the animal introduce blur, obscuring these changes. As a result, a host of behavioral mechanisms, including rapid photoreceptor, retinal, head and body movements have evolved to facilitate image stabilization in sighted animals so as to improve sampling of the image and reduce blur^1,2^. At the same time, in artificial image processing systems, image blur can be reduced in both stationary and moving scenes using a wide range of computational algorithms^3^. However, whether biological systems might utilize computational mechanisms downstream of retinal input to reduce image blur under naturalistic conditions is unknown. Here we show how recurrent circuit feedback dynamically tunes the temporal filtering properties of first order interneurons in the fruit fly *Drosophila* to suppress image blur caused by movement of the animal.

Visual processing in *Drosophila* begins in the retina, where each point in space is sampled by eight photoreceptor cells^4^. Six of these cells, R1-6, are essential inputs for achromatic information processing, including motion detection, and relay their signals to the distal-most region of the optic lobe, the lamina. Within each lamina cartridge, the six R1-6 cells representing the same point in visual space make extensive synaptic connections with three feedforward projection neurons, L1-3. L1-3 then provide differential inputs to downstream neural circuits that are specialized to detect either increases in light intensity or decreases in light intensity^5–7^. Extensive characterization of L1-3 using both artificial and naturalistic stimuli has revealed how each of these cell types can transmit distinct information about the luminance of a scene relative to the local spatial and temporal contrast^8–12^. However, how the temporal dynamics of L1 and L2 might relate to the spatiotemporal structure of the stimulus, particularly over short time intervals, is only incompletely understood.

Photoreceptors depolarize to increases in light intensity and have monophasic impulse responses^13–16^. That is, following a brief flash of light, they transiently depolarize before returning to baseline, a response waveform with a single phase (Figure 1A). The photoreceptor-to-lamina-neuron synapse is inhibitory; therefore, L1-3 hyperpolarize to increases in light intensity and depolarize to decreases in light intensity. Like the photoreceptors, L3 has sustained responses to light and a monophasic impulse response. In contrast, L1 and L2 respond more transiently and have biphasic impulse responses, meaning that they encode information about the rate of change in light intensity, the temporal derivative^17,18^ (Figure 1A). These biphasic impulse responses have been interpreted in the context of efficient coding, a hypothesis in which these neurons maximize the amount of information they encode about the stimulus while minimizing the resources used and remaining robust to noise^17,19–21^. In this framework, contrast changes are preferentially encoded because they contain new, behaviorally salient information. However, the ability of the visual system to estimate the temporal derivative of an input is diminished by blur: as the visual scene moves faster and faster, the measured contrast changes at single points in space diminish. How can neural processing compensate for the effects of blur so as to encode contrast changes if they are blurred in time?

**Figure 1.**
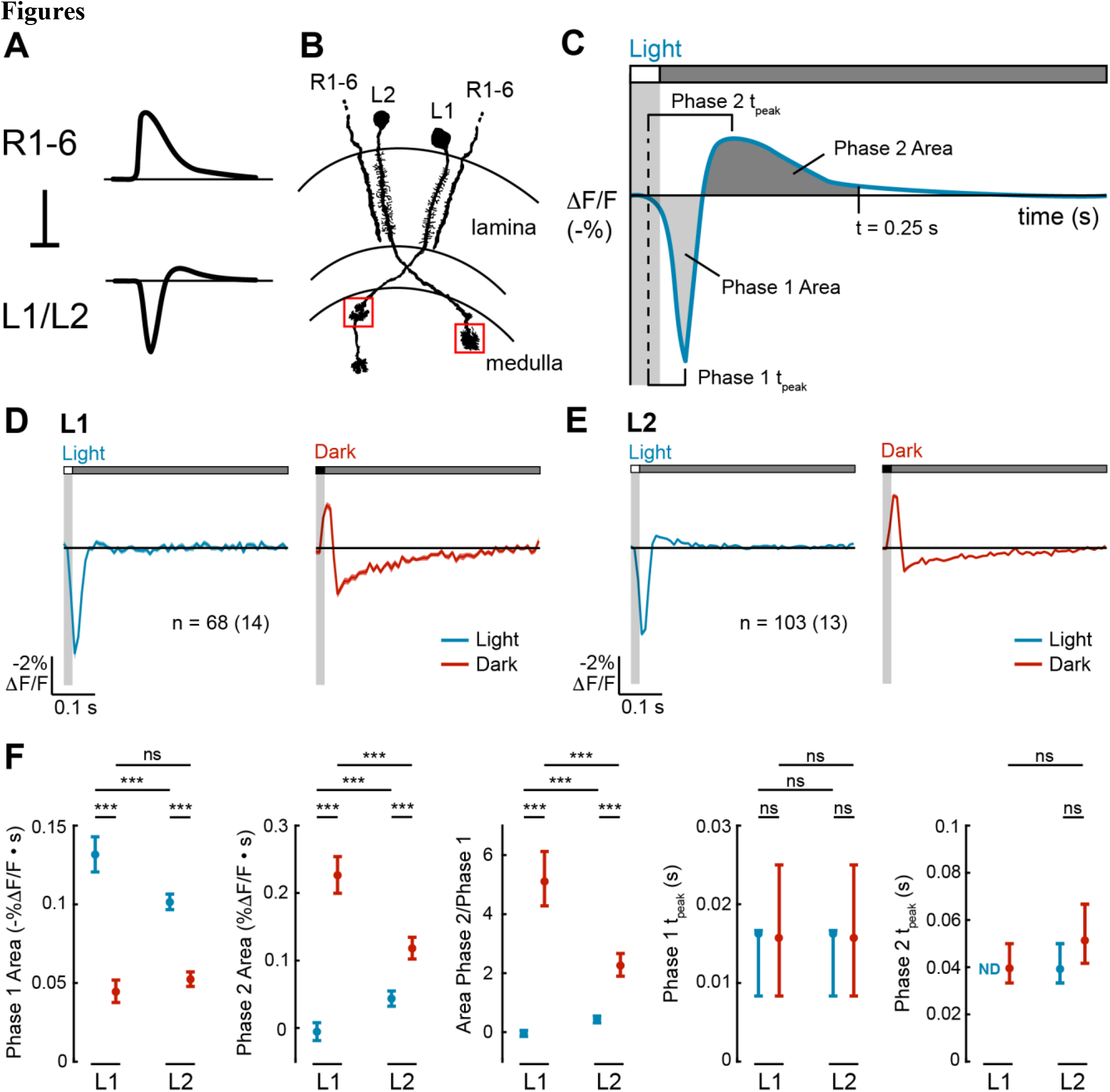
L1 and L2 have biphasic impulse responses that are asymmetric for light and dark. Fig. 1 Image Description (**A)** L1 and L2 are non-spiking neurons that receive inhibitory input from the R1-6 photoreceptors. R1-6 have monophasic responses to light, whereas L1 and L2 have biphasic responses to light. (**B)** L1 and L2 project into the medulla neuropil, where L1 has two arbors (in layers M1 and M5), and L2 has one arbor (in layer M2). We imaged the layer M1 arbor of L1 and the layer M2 arbor of L2 (boxed in red). Diagram adapted from ref^59^. **(C)** Schematic of how the metrics quantifying biphasic impulse responses are defined. The response to a light flash is shown here but equivalent metrics were used to quantify the response to a dark flash. Phase 1 Area: the area under the curve from the time the stimulus-evoked response begins to the time ΔF/F = 0 between the two phases. Phase 2 Area: the area under the curve from the time ΔF/F = 0 between the two phases to 0.25 s after the start of the stimulus. Phase 1 t_peak_: time from when the stimulus-evoked response begins to when it peaks during the first phase. Phase 2 t_peak_: time from when the stimulus-evoked response begins to when it peaks during the second phase. **(D-E)** Responses of L1 (**D**) and L2 (**E**) to a 20-ms light flash (left, blue) or a 20-ms dark flash (right, red) with a 500-ms gray interleave. The solid line denotes the mean response and the shading is ± 1 SEM. n = number of cells (number of flies). (**F)** Quantification of the responses. ND denotes when the Phase 2 Area was 0. The mean and 95% confidence interval are plotted. ns p>0.05, *** p<0.001.

Here we examine temporal processing by L1 and L2 in the fruit fly to address this issue. We find that L1 and L2 impulse responses are biphasic, as observed previously, but are surprisingly nonlinear with respect to light and dark and have larger second phases than expected. Informed by the effect of circuit manipulations on the second phase of L2 responses, we designed a recurrent dynamical model that can recapture the biphasic responses of L1 and L2. This model uses the output of L1 and L2 to scale the strength of a feedback signal back to L1 and L2, creating a recurrent structure. We find that responses that have been shaped by recurrent feedback emphasize rapid transitions between light and dark, producing sharper and faster responses than would be predicted by a model in which the visual system simply tries to reconstruct the “true” contrast change. We show that this sharpening would counteract blur introduced into photoreceptor responses by image motion and by the limited spatial resolution of the eye. Moreover, this recurrent circuitry is tuned by changes in neuromodulatory state that represent movement of the animal and enhances responses to naturalistic stimuli. Thus, temporal processing in early visual interneurons serves not to efficiently encode the signals they receive but instead serves to enhance a critical feature of natural scenes, thereby enhancing downstream extraction of salient visual features.

## Results

### L1 and L2 impulse responses are biphasic and nonlinear with respect to light and dark

We characterized the temporal tuning properties of L1 and L2 by expressing the voltage indicator ASAP2f^22^ cell type-specifically and recording visually evoked signals in the axonal arbors of individual cells (Figure 1B). To do this, we presented 20-millisecond full-field, contrast-matched light and dark flashes off of uniform gray on a screen in front of the fly. These impulse responses represent a linear approximation of how each cell type weighs visual inputs from different points in time and can be quantified with metrics capturing the magnitude and kinetics of each phase of the response (Figure 1C).

In response to the light flash, L1 and L2 both initially hyperpolarized, reaching peak amplitude at the same time (Figures 1D-F). The response of L2 then had a second depolarizing phase, while the response of L1 did not. In response to the dark flash, L1 and L2 initially depolarized, again with identical kinetics, and then hyperpolarized in a second phase. For both cell types, the light and dark impulse responses were not sign-inverted versions of each other, and therefore, these cells were not linear. Instead, the light and dark impulse responses of both cell types were asymmetric, differing in the extent to which they were biphasic. For L1, the light response was monophasic while the dark response was biphasic, with a substantially larger second phase than the first phase (Figures 1D and 1F). For L2, the light and dark responses were both biphasic, but the second phase of the dark response was larger than that of the light response (Figures 1E and 1F). Thus, L1 and L2 process light and dark inputs with distinct kinetics, and strikingly, the second phase of the response was often (but not always) much larger than the first phase of the response.

### Recurrent synaptic feedback and cell-intrinsic properties together shape L2 impulse responses

*Drosophila* photoreceptors have monophasic impulse responses (Figure 1A)^23^, suggesting that the first phase of the response in both L1 and L2 reflects direct photoreceptor input, whereas the second phase of the response originates downstream of photoreceptors. Two non-mutually exclusive categories of biological mechanisms could explain the second phase: circuit elements (such as recurrent synaptic feedback) and cell-intrinsic mechanisms (such as voltage-gated ion channels). To test these possibilities, we took advantage of available genetic tools for cell type-specifically manipulating L2, first by limiting L2 inputs to the photoreceptors alone and then by selectively restoring recurrent feedback driven by L2.

We first measured the impulse responses of L2 in the absence of all circuit inputs other than photoreceptors (Figure 2A). To do this, we blocked synaptic vesicle release by expressing tetanus toxin light chain (TNT) in all neurons directly postsynaptic to photoreceptors using a promoter fragment derived from *ora transientless (ort)*, the histamine-gated chloride channel that serves as the neurotransmitter receptor for photoreceptor input^24–27^. As TNT blocks synaptic transmission instead of altering membrane potential, we could still measure the responses of L2 to photoreceptor input, even as L2 could not relay signals to downstream circuitry. In other words, we isolated L2 from both feedforward and feedback circuits while preserving only photoreceptor input, allowing us to observe how cell-intrinsic mechanisms shape its responses. Silencing circuit inputs to L2 eliminated the second phase of the response to light flashes while increasing the amplitude of the first phase and delaying the time to peak (Figures 2B and 2C). This observation suggests that circuit elements contribute a delayed, depolarizing input that truncates the initial hyperpolarizing response to photoreceptor signals and produces the depolarizing second phase. When we examined the response to dark flashes, we found, to our surprise, that blocking circuit input did not eliminate the second phase (Figures 2B and 2C). Instead, the second phase became larger and more sustained, peaking at a later time. The first phase was also unchanged, demonstrating that photoreceptor input and cell-intrinsic mechanisms are sufficient to produce it. Thus, the biphasic character of the light impulse response of L2 requires circuit input, while that of the dark impulse response is determined largely by cell-intrinsic mechanisms.

**Figure 2.**
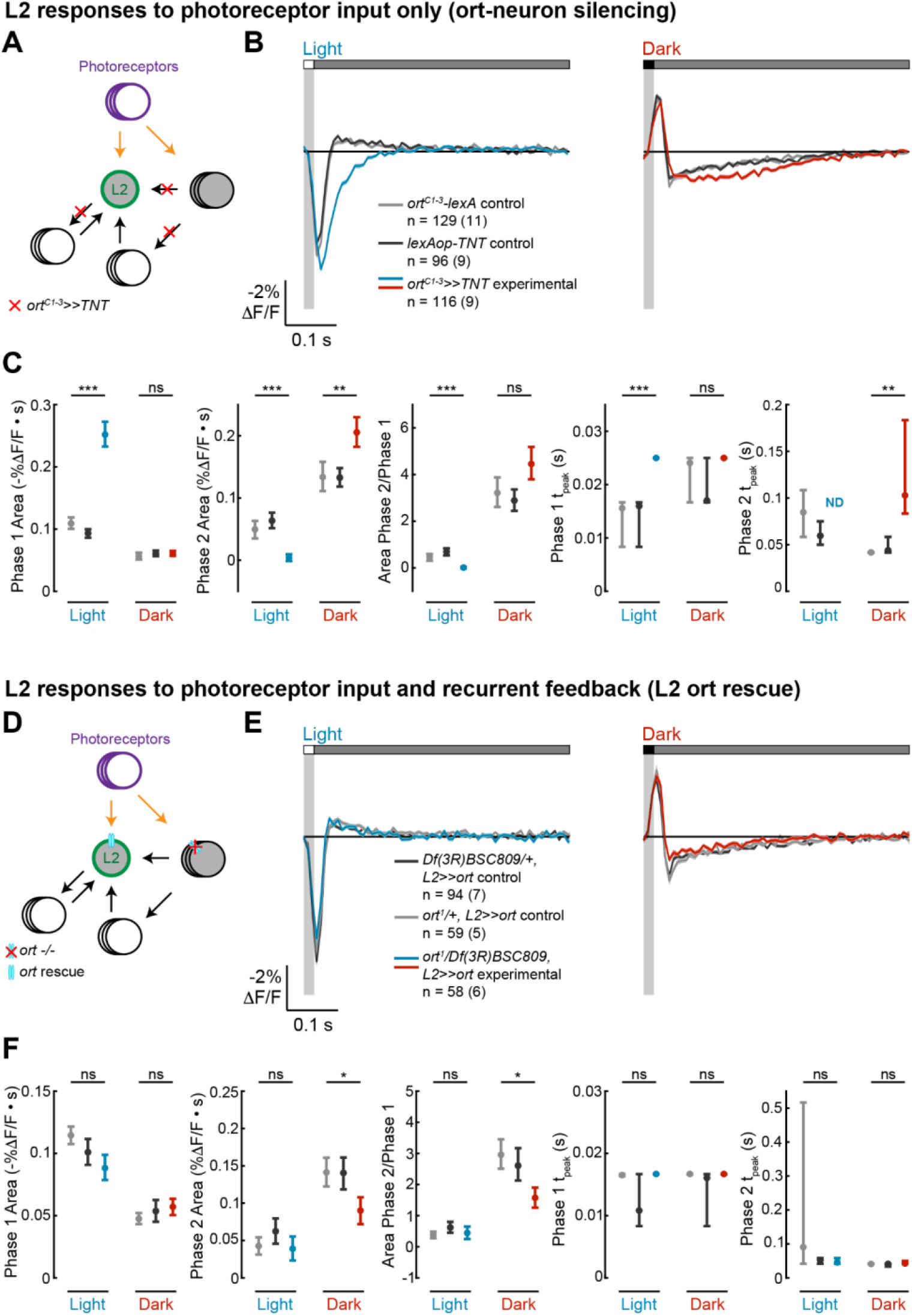
Circuit and intrinsic mechanisms shape L2 impulse responses. Fig. 2 Image Description **(A-C)**, ort-neuron silencing. **(D-F)**, L2 ort rescue. **(A)**, Diagram of the ort-neuron silencing experiment: *ort*^*C1-3*^*-lexA*-driven tetanus toxin (TNT) blocks synaptic vesicle release from all neurons that receive direct input from photoreceptors, thereby preventing visual information from being transmitted through circuits downstream of the first-order interneurons. **(D)** Diagram of the L2 ort rescue experiment: in an *ort* null background, *ort* is rescued specifically in all L2 neurons. This preserves L2 responses to photoreceptor input and any L2-dependent circuit feedback but eliminates any circuit-based visual input that does not pass through L2. **(B, E)** Responses of L2 to a 20-ms light flash (left) or a 20-ms dark flash (right) with a 500-ms gray interleave. The responses of the genetic controls are in gray; the experimental responses are in color. The solid line is the mean response and the shading is ± 1 SEM. n = number of cells (number of flies). **(C, F)** Quantification of the responses in (**B, E**). The mean and 95% confidence interval are plotted. ND is shown instead of the Phase 2 t_peak_ value when the Phase 2 Area is 0. ns p>0.05, * p<0.05, ** p<0.01, *** p<0.001, Bonferroni correction was used for multiple comparisons.

We next performed a complementary experiment in which only L2 could respond to light and activate downstream circuitry. We leveraged the fact that *ort*, a histamine gated chloride channel, is essential for photoreceptors to signal to neurons in the lamina^24,25,28,29^. As expected, L2 neurons in *ort*-null flies did not respond to light (Figure S1). Next, we rescued expression of *ort* specifically in L2 in an *ort*-null genetic background, thereby allowing L2 alone to receive photoreceptor input and signal to its postsynaptic partners. (Figure 2D). Thus, circuit contributions that depend on direct visual input to neurons other than L2 were blocked, but L2-dependent recurrent circuits and cell-intrinsic mechanisms were preserved. Strikingly, L2-specific *ort* rescue restored not only the initial, photoreceptor-mediated phase but resulted in biphasic impulse responses to both light and dark flashes (Figures 2E and 2F). The light impulse response was indistinguishable from those measured in control flies heterozygous for each of the *ort* null alleles, indicating that L2 function is sufficient to produce the biphasic response and that the circuit mechanisms that generate the depolarizing second phase are L2-dependent and hence recurrent. However, interestingly, while the dark impulse response was biphasic, the rescue was not complete: the first phase of the response was intact, but the amplitude of the second phase was smaller than that seen in control animals. Therefore, L2-independent circuit mechanisms contribute to this second phase, consistent with the results from the *ort-TNT* experiments (Figure 2B). Taken together, these results demonstrate that both cell-intrinsic properties and L2-driven recurrent feedback shape impulse responses in L2 in an asymmetric, stimulus-dependent fashion.

### A recurrent model replicating L1 and L2 response dynamics counteracts motion-induced blurring

Given the stimulus-selective biphasic impulse responses (Figure 1) and their complex control mechanisms (Figure 2), we next examined how the temporal properties of L1 and L2 affect visual processing. Biphasic impulse responses are generally thought of as biological implementations of derivative-taking, or in the frequency domain, bandpass filters. In this framework, the biphasic waveform that best approximates derivative-taking—or equivalently, is most strongly bandpass—has first and second phases with identical shapes and integrated areas. However, we never observed an impulse response of this form (Figures 1D-1F). Instead, the second phase of the response was often much stronger than the first phase (Figures 1D-1F), a form that has not been observed in many other visual neurons^16,23,30–35^. A linear filter with that shape would be low-pass in the frequency domain and would predict responses to contrast steps that are inconsistent with physiological measurements (Figure S2)^22,36^. Furthermore, as noted above, the observation that the light and dark impulse responses are not sign-inverted versions of each other indicates that temporal processing by L1 and L2 is not linear (Figures 1D-1F).

To explore the functional consequences of these properties, we modeled L1 and L2 responses using a set of differential equations that describe the time-varying activity of each cell (*v*) as a function of feedforward input from photoreceptors as well as feedback from recurrent circuitry (Figure 3A). The feedback (*y*) was modeled as a longer time-constant, sign-inverted signal scaled by the output of each cell type and served to generate the second phase of the L1 (or L2) response. One parameter (*g*) controlled how strongly depolarization or hyperpolarization of L1 (or L2) engages downstream feedback circuits, while a second parameter (*w*) controlled the overall strength of feedback. This recurrent, dynamical model captured the biphasic impulse responses of L1 and L2, including the nonlinear, asymmetric second-phase responses to light and dark flashes (Figure 3B). Moreover, different sets of model parameters could reproduce the differences between the two cell types (Figure 3B and Methods).

**Figure 3.**
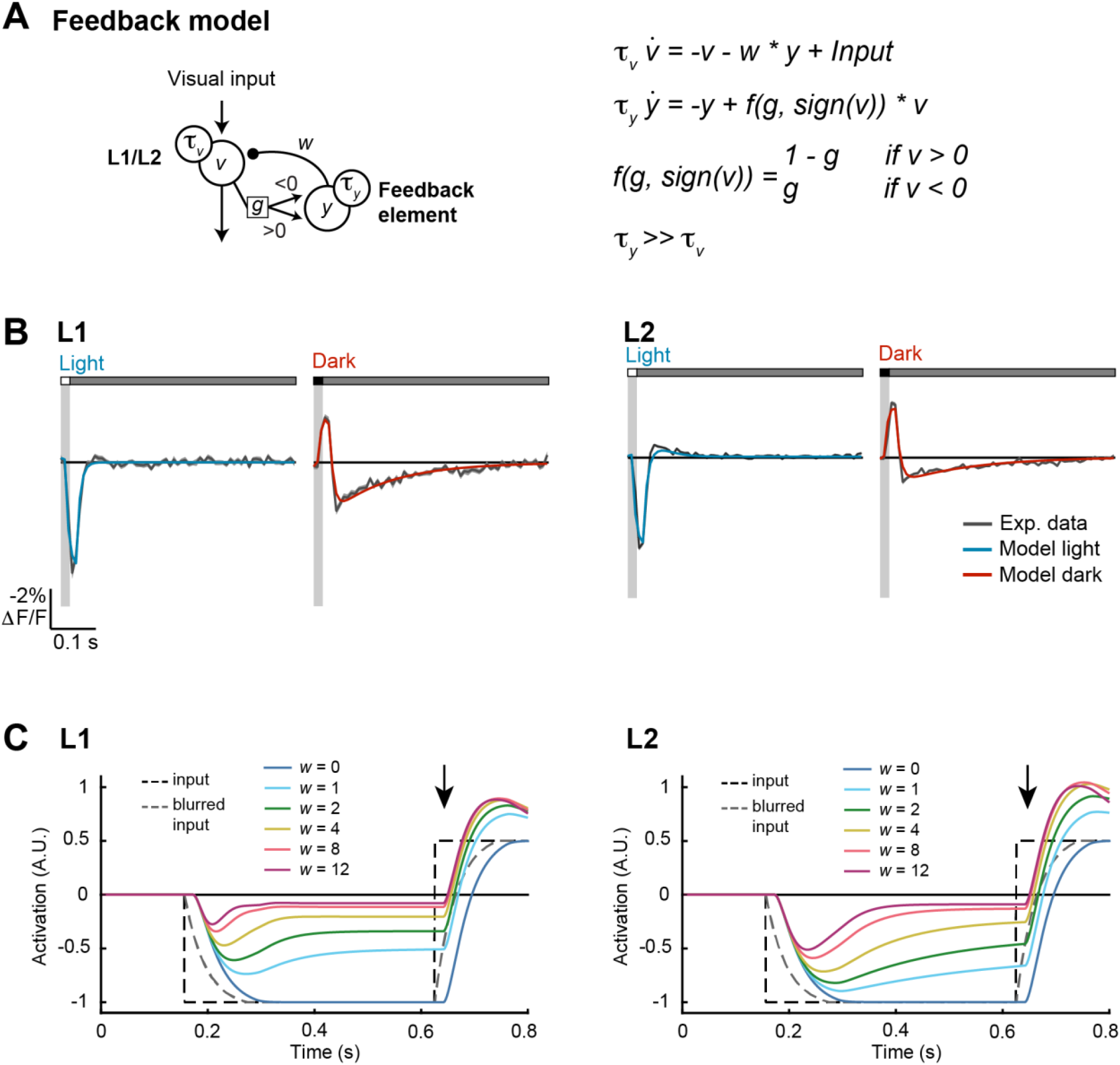
A recurrent feedback model recapitulates L1 and L2 responses and deblurs moving edges. Fig. 3 Image Description **(A)** Schematic of the feedback model (left) and equations (right). *v* is the neural response of L1/L2, and 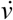 is its time derivative. *y* is the output of the feedback element, and 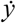 is its time derivative. τ_*v*_ and τ_*y*_ describe time constants for *v* and *y*, respectively. *w* is the strength of feedback, and *g* controls how strongly positive and negative values of *v* engage feedback. **(B)** Modeled L1 (left) and L2 (right) responses to 20-ms light (blue) or dark (red) flashes overlaid on experimentally measured responses of L1 and L2 from Figure 1D (gray). **(C)** Modeled L1 (left) and L2 (right) responses to a motion-blurred edge (blurred input, gray dashed line; original input, black dashed line) for models with different feedback weights (*w*). No feedback is *w* = 0 (dark blue). As the weight increases, the model responds more rapidly to the dark-to-light transition (see responses at the arrow). Responses are plotted with inversion over the x-axis to facilitate comparisons with the inputs.

In this model, the sign of the feedback and its long time constant can generate responses that are of opposite sign and perdure beyond the initial response. For example, a dark flash produces depolarization followed by sustained hyperpolarization. As the initial response to a light flash is a hyperpolarization, the second phase of the dark response could transiently enhance subsequent responses to light. We therefore reasoned that this property would be particularly impactful for stimuli that contain sequences of opposite contrast polarities. Such stimuli are common with motion; for example, a dark object passing over a light background will produce a temporal sequence of dark followed by light.

As a test case, we examined how the model responded to simple artificial stimuli corresponding to dark or light steps (Figure 3C). By definition, the transition in the stimulus was instantaneous; however, the input to the modeled L1 or L2 was delayed and blurred in time, reflecting the limited temporal resolution of photoreceptors and the upstream synapse. We then varied the weight of the feedback signal; when *w* = 0, the modeled L1 or L2 responses followed the time course of this delayed and blurred input. However, as the weight of the feedback increased, the transition reported by the modeled response became faster and sharper, more closely approaching the original instantaneous transition. The degree to which the model could accelerate responses to dark or light inputs depended on the degree of asymmetry in the strength of the feedback (Figure S3). In particular, when g = 0, meaning that only dark inputs drive feedback, responses to transitions that follow a dark input were selectively sharpened; in contrast, when g = 1, meaning that only light inputs drive feedback, responses to transitions that follow a light input were selectively sharpened. When g = 0.5, transitions in both directions were sharpened. In this context, L1 had highly asymmetric feedback (g = 0.0277) and accordingly, edge enhancement occurred mainly for one polarity (Figure S4). Conversely, L2 was also asymmetric, but less so (g = 0.169), and enhanced edges of both polarities. Notably these effects on temporal sharpening would be preserved by the known contrast selectivities of downstream neurons (Figure S4). Thus, recurrent feedback generates a form of temporal edge enhancement that could be useful for downstream processing.

### Recurrent feedback compensates for motion-induced blurring of natural images

An animal’s own movement blurs stationary visual inputs by reducing their apparent contrast in both time and space and by imposing a temporal delay between the true and perceived location of objects in the environment. Given the dynamical model’s ability to sharpen an artificial moving edge (Figure 3C), we hypothesized that temporal sharpening could also reduce blur generated by self motion, leading to responses that are closer in time to the original, non-blurred signal. To test this, we simulated the responses of L1 and L2 evoked by moving natural images^37,38^. In brief, the natural images were processed with spatial blurring corresponding to retinal optics and temporal blurring produced by the non-zero integration time of photoreceptors and the effects of self-motion (Figure 4A and Methods). The blurred images were then used as inputs to the dynamical model. For comparison, we also used these images as inputs to a model that lacked recurrent feedback (by setting *w* = 0). The outputs of these models were then compared with the input images to determine the temporal offset that maximized spatial alignment (Figure 4B). This offset captures the delay caused by both motion blur and circuit processing. Strikingly, the model that incorporated recurrent feedback significantly reduced the temporal offset relative to the model without feedback (Figure 4B). Intuitively, this effect arises because the temporal shift induced by feedback in the recurrent circuit compensates for the temporal shift induced by motion, thereby deblurring the image.

**Figure 4.**
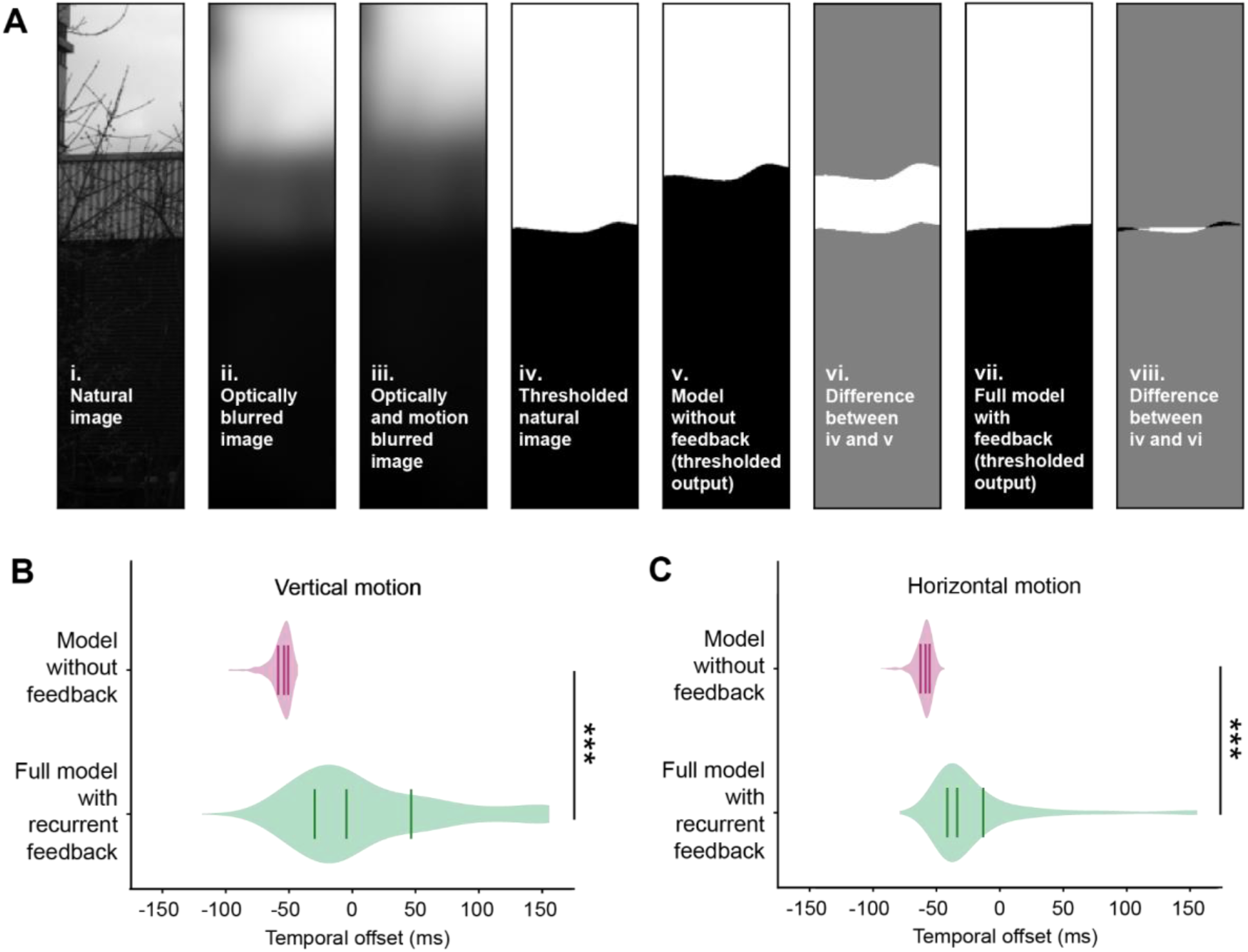
A feedback model deblurs and strengthens responses to edges. Fig. 4 Image Description **(A)** Deblurring edges in natural images. Thresholding refers to binarizing the image according to contrast polarity (above zero: white, below zero: black). **(B)** Violin plots of the temporal offset between the thresholded natural image (**A-iv**) and either the thresholded output of the model without feedback (**A-v**) or the full model with recurrent feedback (**A-vii**), for 464 natural images (see Methods). Images were blurred with motion in the vertical direction. Lines indicate the lower quartile, median, and upper quartile. *** p<0.001, Wilcoxon signed-rank test. **(C)** As in **(B)**, but with motion in the horizontal direction.

### Octopaminergic neuromodulation enhances feedback and improves deblurring at fast speeds

The average speed of images across the retina differs greatly depending on whether a fly is stationary, walking, or flying–thereby varying the motion blur. As a result, behavioral context should change the temporal processing strategy of L1 and L2. Indeed, many visual neurons downstream of L1 and L2 have different temporal tuning in moving and quiescent flies^35,39–44^. Using our model, we asked how changing the strength of the feedback element changes responses to high speed and low speed images. We found that increasing feedback improved deblurring of fast-moving images, diminishing the temporal offsets relative to a model with no feedback (Figures 5A and S5). Critically, increasing feedback also worsened blurring of slow-moving images, leading to “over-corrected” temporal offsets that frequently exceeded 0 ms, particularly for motion in the vertical direction (Figure 5A). Thus, to be most useful, feedback should adapt to different contexts. Specifically, feedback should be stronger in behavioral states where visual image motion is faster, such as flight. To test this model prediction, we applied the octopamine agonist chlordimeform (CDM) to the brain, a manipulation that has been used to mimic the change in neuromodulatory state associated with flight and that enhances tuning for faster image velocities in visual neurons^35,40,43^. Under these conditions, the measured impulse responses of L1 and L2 to dark flashes had larger second phases relative to their first phases, while their light responses were relatively unchanged, qualitatively matching predictions of the model with enhanced feedback (Figures 5B and 5C). Thus, temporal filtering by L1 and L2 can be shaped by octopamine to enhance deblurring under behaviorally relevant contexts.

**Figure 5.**
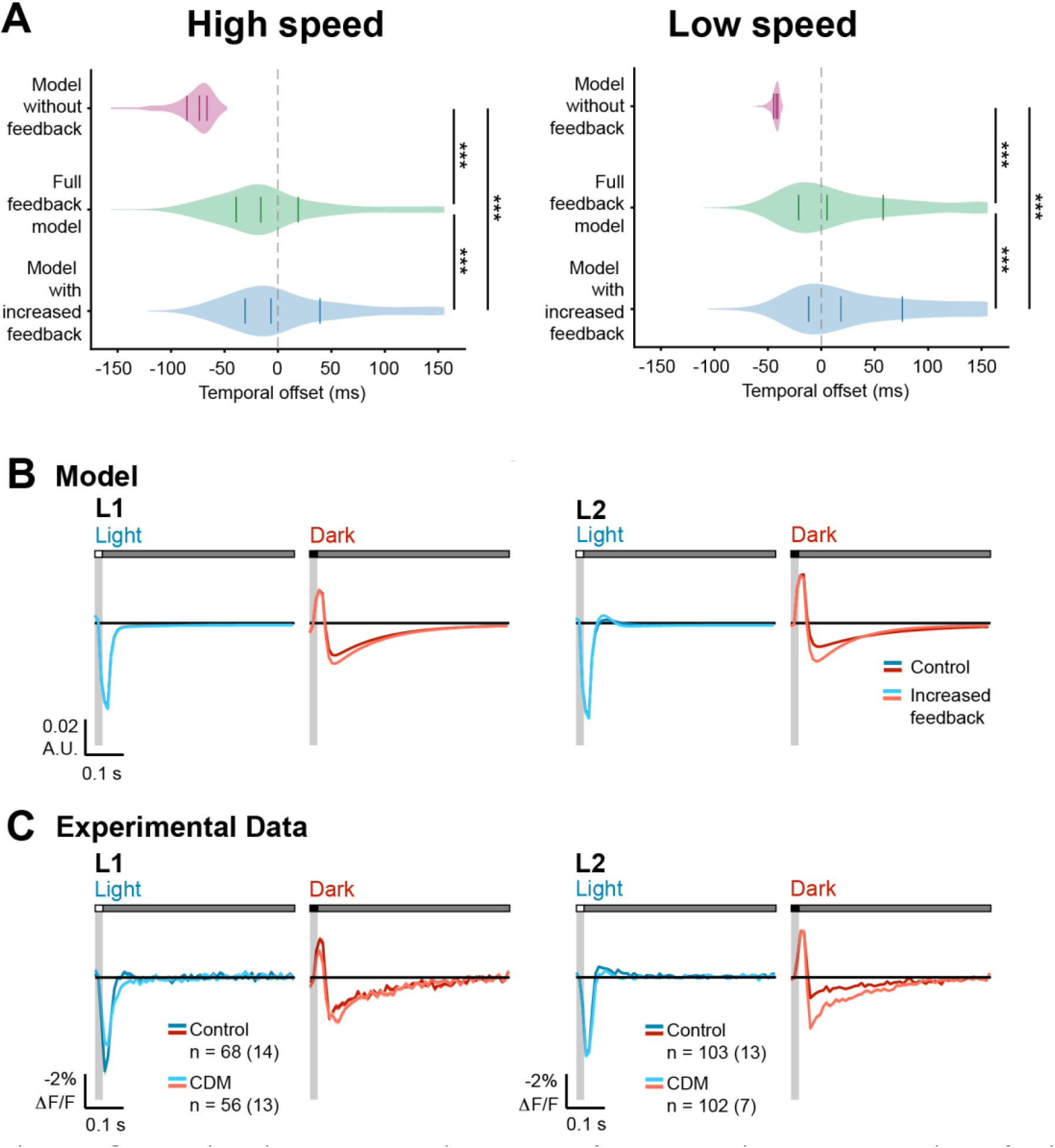
Octopaminergic neuromodulation enhances feedback and improves deblurring at fast image speeds. Fig. 5 Image Description **(A)** Violin plots of the temporal offset between the natural image and model output at high (left, 600 °/s) and low (right, 150 °/s) image speeds, using *w* = 0 (model without feedback), *w* = 12.7 (full feedback model), and *w* = 24 (model with increased feedback). Images were blurred with motion in the vertical direction. Solid lines indicate the lower quartile, median, and upper quartile. Dotted gray line indicates temporal offset = 0 ms. *** p<0.001, Wilcoxon signed-rank test. **(B)** Modeled L1 (left) and L2 (right) responses to 20-ms light (blue) or dark (red) flashes with a 500-ms gray interleave under control or enhanced feedback. **(C)** Experimentally measured responses of L1 and L2 to a 20-ms light flash (blue) or a 20-ms dark flash (red) with or without bath application of the octopamine receptor agonist CDM (20 μM). The control data is repeated from Figure 1. The solid line is the mean response and the shading is ± 1 SEM. n = number of cells (number of flies).

### Feedback sharpens intensity changes in naturalistic stimuli over time

To test the generalizability of these predictions, we recorded the voltage responses of L2 to a temporally naturalistic stimulus. This stimulus simulated the sequence of light intensities that would be detected by the photoreceptors upstream of a single L2 cell as a fly moves relative to the visual scene (Figures 6A, S6A, and S6B). Under these conditions, the voltage responses of L2 broadly followed the changes in stimulus intensity, depolarizing as the stimulus became dimmer and hyperpolarizing as the stimulus became brighter (Figure 6B). Interestingly, responses to small contrast changes were accentuated and responses to large contrast changes were accelerated (Figure 6B, insets), resulting in temporally sharpened transitions.

**Figure 6.**
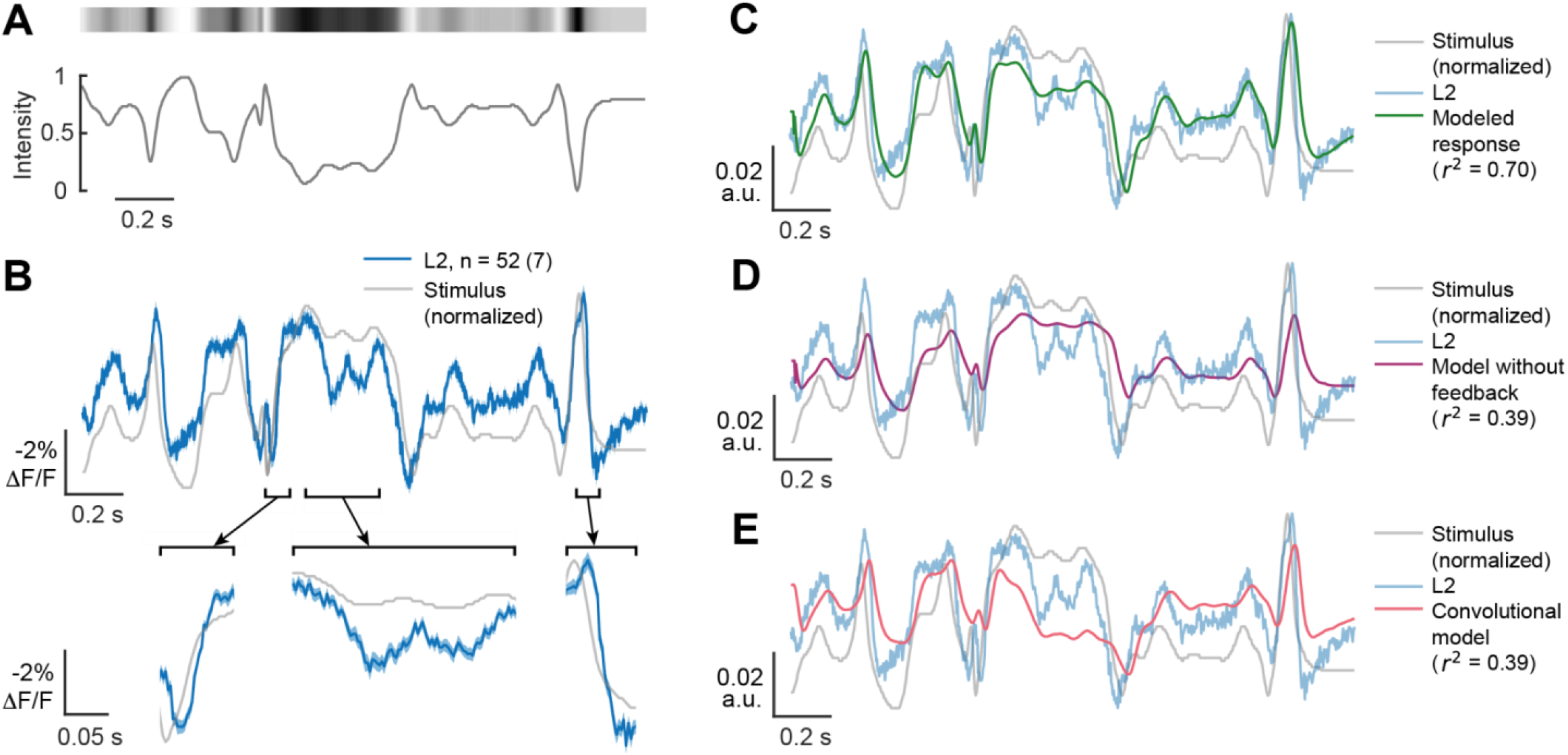
Both measured and modeled responses sharpen responses to naturalistic stimuli. Fig. 6 Image Description **(A)**. The time varying changes in intensity of a full-field naturalistic stimulus (bottom) and a visual representation of the stimulus intensity (top). **(B)** The measured response of L2 (blue, top) to this naturalistic stimulus, overlaid on an inverted stimulus trace (gray). Selected segments are depicted below as expanded insets (bottom) to better visualize response dynamics. The solid line is the mean response and the shading is ± 1 SEM. n = number of cells (number of flies). **(C)** Full feedback model of L2 responses to the naturalistic stimulus using the same parameters as for the impulse responses, overlaid on the inverted stimulus (gray) and the measured response (light blue). **(D)** As in **(C)**, but using the model without feedback (magenta). **(E)** As in **(C)**, but with a convolutional model using the impulse response as a linear filter (red).

We then modeled L2 responses to the naturalistic stimulus using the same parameters that were used to predict the impulse response. Despite the distinct stimulus statistics, this recurrent model predicted the response of L2 to the naturalistic stimulus well (r^2^ = 0.72) (Figure 6C). Modifying the model to remove the feedback component or using a convolutional model that used the impulse response as a linear filter dramatically reduced performance (r^2^ = 0.39 for both) (Figures 6D and 6E). Directly fitting the model to the naturalistic stimulus response improved performance (r^2^ = 0.90), while using the same model but without feedback only yielded r^2^ = 0.56 (Figure S6). Taken together, these measured and modeled results suggest that the recurrent structure of the dynamical model captures the temporal features of L2 responses under naturalistic conditions and further support a role for L2 in deblurring stimuli.

## Discussion

Responding strongly and precisely to changes in contrast is a critical step in visual processing. Using *in vivo*, two-photon voltage imaging and computational modeling to measure and describe the temporal properties of the first order interneurons L1 and L2, we reveal surprising complexity in this computation. Examining the impulse responses of these cells, we show that L1 and L2 have distinct responses to flashes of light and dark. Across both cell types, the responses to these oppositely signed flashes differ in waveform and thus are not consistent with a linear system. Intriguingly, we observed that the response to dark flashes had an unusual biphasic character, where the second phase of the response was much larger than the first. Using genetic manipulations that isolated different elements of the circuit, we show that while cell intrinsic mechanisms are important, recurrent circuit feedback plays a central role in shaping the biphasic responses of L2 to both light and dark. To test hypotheses about the functional roles of these responses, we built a dynamical model with recurrent feedback that was fit to experimental data. Using simulation, we find that the recurrent feedback element of the model enables faster responses to contrast changes, compensating for motion-induced blurring of natural scenes. This feedback can be tuned by modulating octopaminergic signaling, consistent with adapting the feedback to the expected degree of blur imposed by self-motion. Finally, we show that L2 responds sharply to contrast changes in a naturalistic stimulus, as does the recurrent feedback model, even when fit to the impulse responses. Taken together, tunable recurrent feedback early in visual processing represents a computational mechanism that can compensate for spatiotemporal blurring.

Our study builds upon previous work extensively characterizing the role of L1 and L2 in relaying information from R1-6 photoreceptors to downstream circuits. Prior work has shown that L1 and L2 encode information about contrast and provide inputs to downstream circuits that become selective to light and dark^5,6,8,9,22,33,45–48^. In addition to encoding contrast, L1 and L2 also adapt to recent stimulus history, rescaling their responses relative to the frequency and luminance distributions of the input over a timescale of hundreds of milliseconds to seconds^49^. Genetic and circuit dissection experiments have suggested that the response properties of L1 and L2 are not solely dependent on photoreceptor input, but rather are a product of dynamic lateral interactions among multiple cell types, as well as recurrent feedback connections with photoreceptors^36,50–52^. We have built upon these previous findings by taking temporally precise, cell type-specific voltage measurements of L1 and L2 axon terminals in response to both artificial and naturalistic stimuli. As expected, the responses of L1 and L2 to these visual stimuli are consistent with contrast tuning. Interestingly, these contrast tuned responses were unusually sharp in time, particularly in response to the naturalistic stimulus at times when the stimulus contrast changed abruptly, creating a temporal “edge”. Using dynamical modeling informed by circuit perturbations, we show that this phenomenon emerges from oppositely signed recurrent feedback. Intuitively, this effect arises because if the stimulus maintains one intensity and then transitions to a different intensity, the recurrent input’s slower time constant causes a normally dampening input to become transiently facilitating, accelerating responses to large contrast changes. This sharpening effect emerges over timescales on the order of 100 milliseconds, a timescale over which changes commonly occur in moving natural stimuli.

Predictive coding provides one explanation for the careful temporal shaping seen in L1 and L2 by positing that a goal of early sensory processing is to efficiently encode sensory information^17,19^. In this view, temporal impulse responses are biphasic so as to encode information only about changes in a stimulus, minimizing the use of neural resources when inputs are constant and therefore predictable. However, this assumes that the main goal of the initial steps in sensory processing is to recode the input ensemble in a more efficient manner. Rather, nervous systems could recode the information in a way that is not necessarily veridical with respect to the information in the stimulus but instead in a way that would make it easier to perform downstream computations useful for behavior^53,54^.

In parallel, control theory in engineering stresses the ability of feedback to change the output dynamics of a system that has constrained input components^55^. We interpret our findings in this light. Feeding back a delayed, scaled, and sign-inverted version of a sensory neuron’s input through a recurrent circuit combined with carefully tuned intrinsic properties makes the system hypersensitive to input changes. This can create responses that occur earlier and are disproportionately larger than specified by the sensory input. This strategy is useful in that it can not only undo blurring of edges during self-motion but can also overemphasize them. In other words, it exaggerates critical features, creating a caricature-like representation of the sensory input to feed into downstream computations. Importantly, the strength of the feedback is adjustable, thereby allowing deblurring to be matched to the expected characteristics of the input. Thus, in this view, even the most peripheral visual circuits in the fly implement critical processing steps that enhance specific visual features.

This type of tunable temporal sharpening could be applied to many sensory systems. In vertebrates, bipolar cells, as the first neurons downstream of the photoreceptors, are the circuit analogs of L1 and L2. However, while the linear filters of many bipolar cell types are biphasic, the second phase is the same size or smaller than the first phase and is not extended in time, unlike L1 and L2^32,56,57^. We speculate that this difference between *Drosophila* and vertebrates is because phototransduction is faster in flies, perhaps better preserving the contrast changes to be sharpened. Conversely, auditory processing in vertebrates can detect rapid changes in stimulus features and use this information to guide behavior. Remarkably, in echolocating bats, neurons in inferior colliculus display biphasic responses in which the second phase is extended in time, a receptive field structure that enhances contrast along multiple stimulus axes^58^. Thus, in sensory systems where detecting rapid stimulus changes is critical, this form of temporal deblurring might provide a generalizable strategy for beginning to abstract the complexity of the natural world into a set of enhanced features primed for downstream computations.

## Acknowledgements

Work done at Stanford is performed on unceded land of the Muwekma Ohlone Tribe. We thank Chi-Hon Lee for sharing *21D-GAL4, ort*^*1-3*^*-lexA, LexAop-TNT*, and *UAS-ort* flies. We thank Bella Brezovec for collecting the fly walking data used to generate the naturalistic stimulus and members of the Clandinin lab for feedback and troubleshooting assistance. This work was supported by R01EY022638 (T.R.C.), by a National Defense Science and Engineering Graduate (NDSEG) Fellowship (M.M.P.), a Stanford Interdisciplinary Graduate Fellowship (H.H.Y.), a Jane Coffin Childs Fellowship (H.H.Y.), an NIH K99 (NS129759 to H.H.Y.), R01EB028171 (F.C. and S.D.), and U19NS104655 (S.D. and T.R.C.). T.R.C. is a Chan-Zuckerberg Biohub Investigator.

## Author Contributions

Conceptualization, T.R.C. and H.H.Y.; Methodology, M.M.P., F.C., M.X., S.D., and H.H.Y.; Formal Analysis, M.M.P., F.C., M.X., S.D., and H.H.Y.; Investigation, M.M.P, M.X., and H.H.Y.; Writing – Original Draft, M.M.P., T.R.C., and H.H.Y.; Writing – Review and Editing, M.M.P., F.C., M.X., S.D., T.R.C., and H.H.Y.; Visualization, M.M.P., F.C., and H.H.Y.; Supervision, S.D. and T.R.C.; Funding Acquisition, S.D. and T.R.C.

## Declaration of Interests

The authors declare no competing interests.

**Figures**

## Supplemental Figures

**Figure S1.**
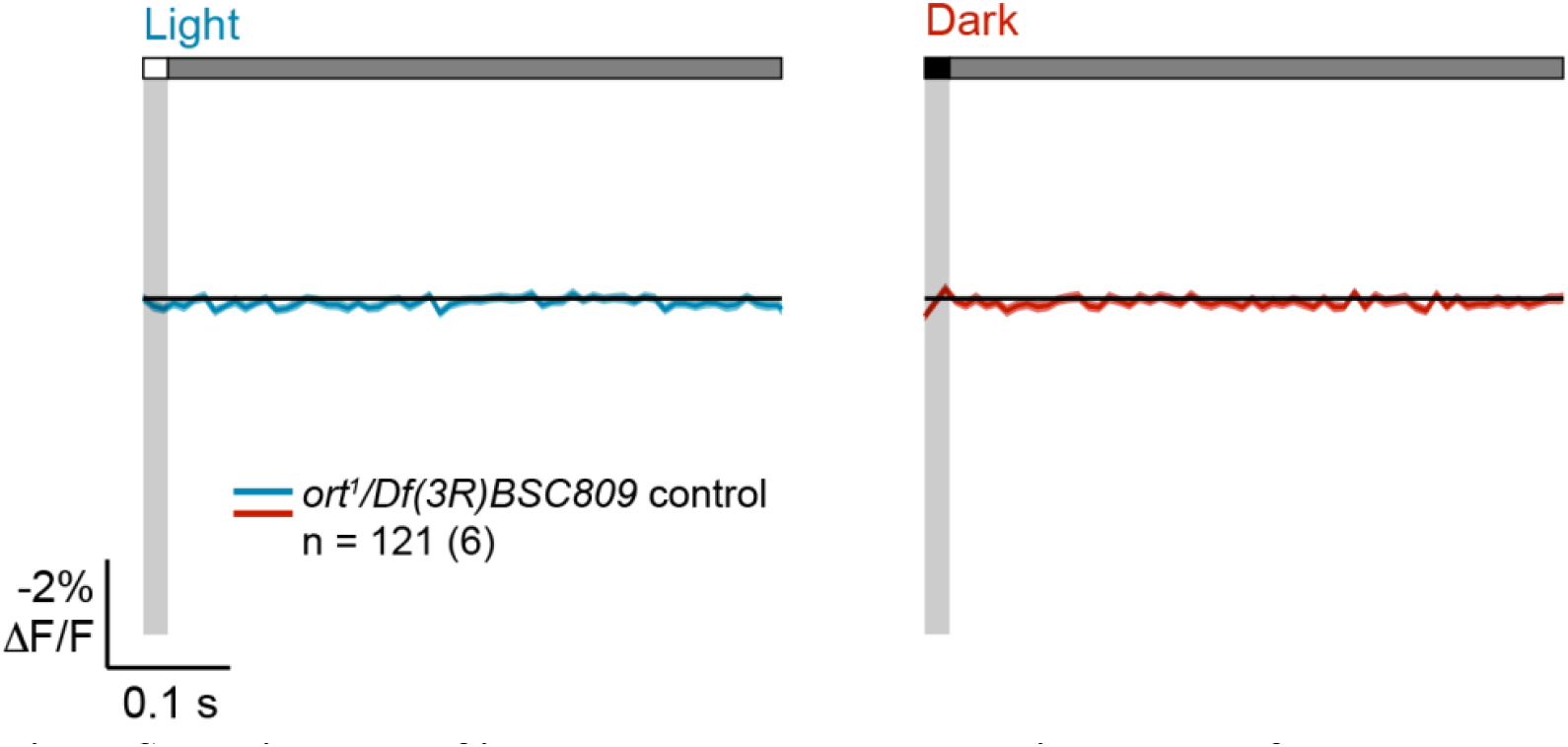
L2 in *ort-*null flies does not respond to 20-ms light or dark flashes. Fig. S1 Image Description Responses of L2 in *ort*^*1*^*/Df(3R)BSC809* flies lacking *ort* rescue to a 20-ms light flash (left, blue) or a 20-ms dark flash (right, red) with a 500-ms gray interleave. The solid line is the mean response and the shading is ± 1 SEM. n = number of cells (number of flies).

**Figure S2.**
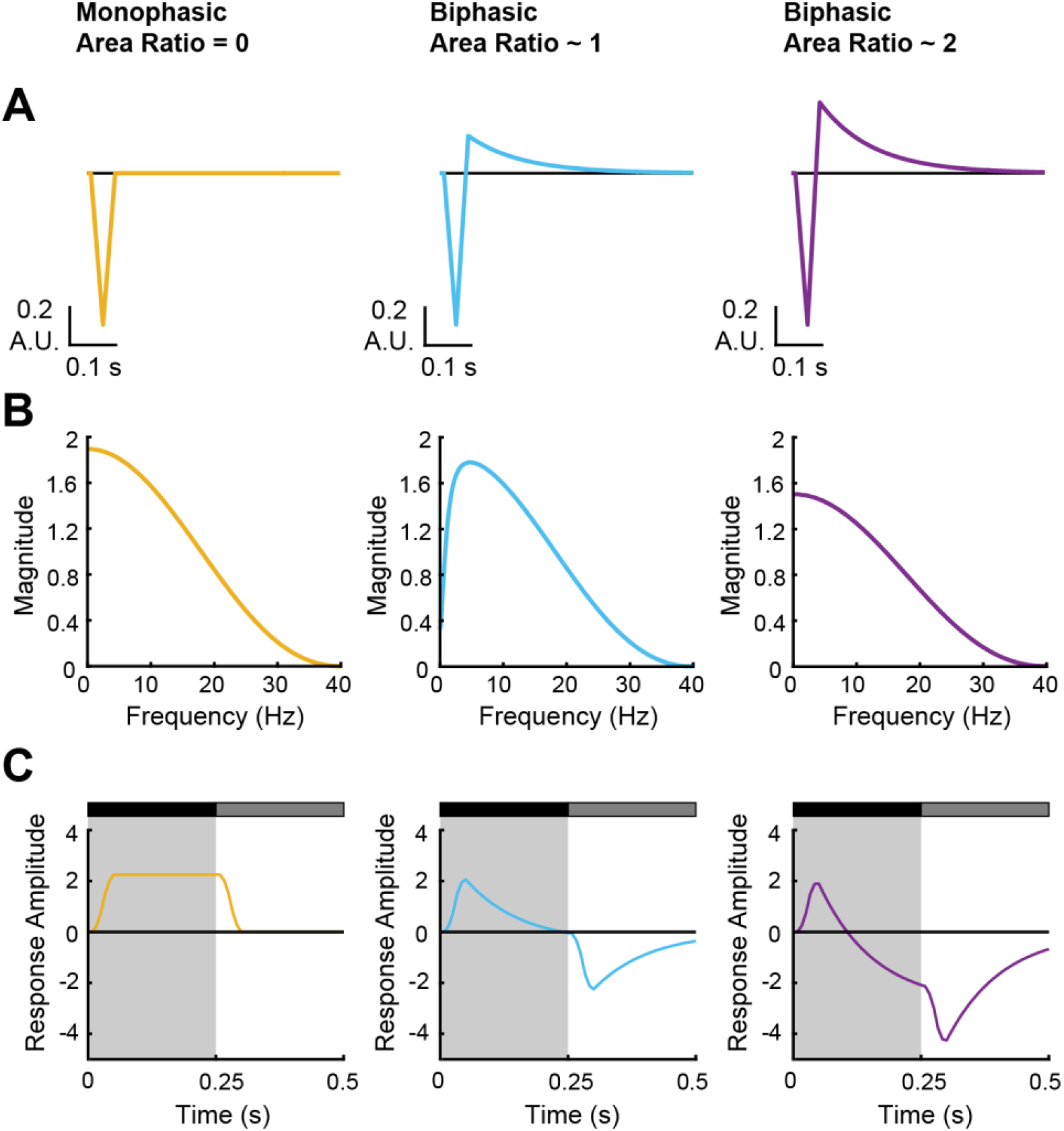
Linear filtering properties of idealized impulse responses. Fig. S2 Image Description **(A)** Idealized impulse responses that approximately match those measured throughout this study. The peak amplitude of the second phase was varied to produce monophasic or biphasic responses with a specific ratio between the areas of the first and second phases; the time-to-peak of the first and second phases as well as the decay of the second phase were held constant. Response amplitude is in arbitrary units (A.U.). **(B)** The magnitude across frequencies for each of the impulse responses in (**A**), computed as the absolute value of the Fourier transform. Note that when the second phase becomes larger than the first phase (area ratio = 2), the filter is low-pass like the monophasic impulse response. **(C)** Simulated response to a 250-ms dark step off of gray computed by convolving the impulse responses in (**A**) with the stimulus. Note the delayed hyperpolarization produced by the area ratio = 2 impulse response.

**Figure S3.**
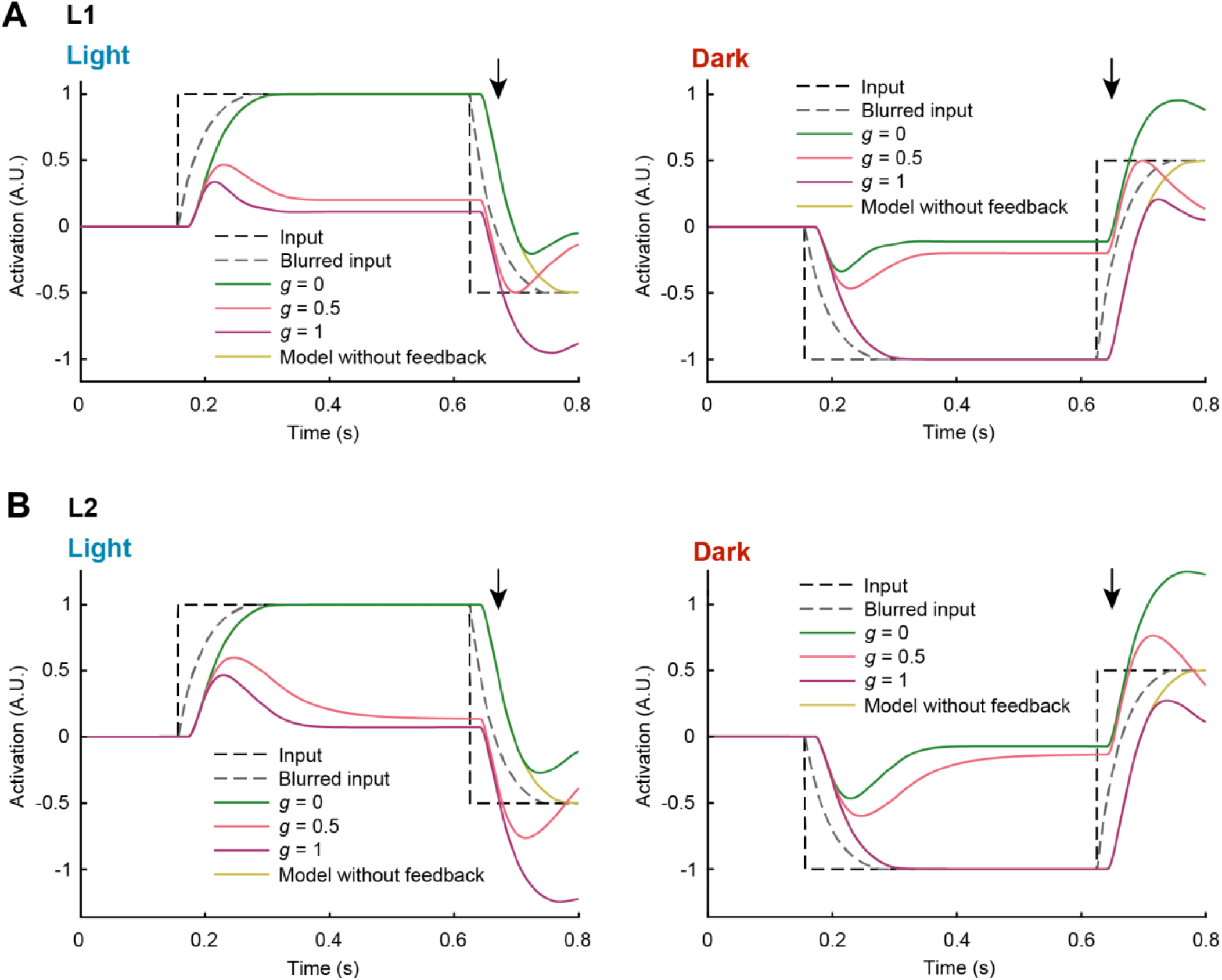
Asymmetric adaptation differentially deblurs and enhances light and dark edges. Fig. S3 Image Description **(A)** Simulated responses to a motion-blurred light edge (left) and dark edge (right) for models with different degrees of asymmetric adaptation (g values). All other parameters are the same as those used to simulate L1 impulse responses in Figure 3. When *g* = 0, only dark drives the feedback element; when *g* = 1, only light drives the feedback element; when *g* = 0.5, both light and dark equally drive the feedback element. Note that when *g* = 0, the response following a dark-to-light transition is faster and stronger, but the response to a light-to-dark transition is like that of the no-feedback model. When *g* = 1, the response following a light-to-dark transition is faster and stronger, but the response to a dark-to-light transition is like that of the no-feedback model. When *g* = 0.5, both light and dark edges are deblurred and enhanced, but not as strongly as when *g* = 0 or 1. Responses are plotted with inversion over the x-axis to facilitate comparisons with the inputs. **(B)** As in **(A)**, but with parameters used to simulate L2 impulse responses in Figure 3.

**Figure S4.**
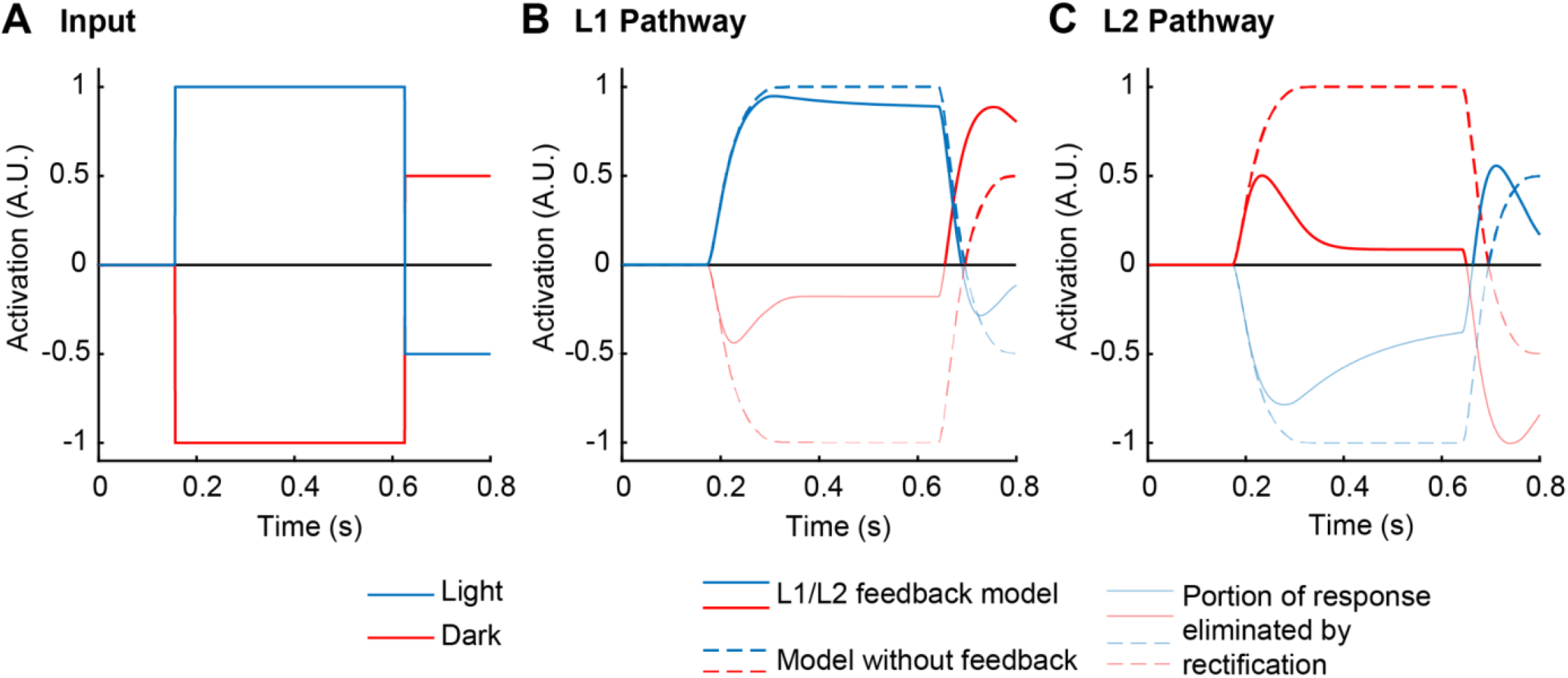
Half-wave rectification preserves deblurring and enhancement of edges in the L1 and L2 pathways. Fig. S4 Image Description **(A)** Light (blue) and dark (red) edge inputs. **(B)** Simulated responses to motion-blurred light and dark edges by an L1 pathway-like model incorporating L1-type dynamic feedback followed by a sign-inversion and half-wave rectification. Dotted lines are the responses of a no-feedback model. Thin lines are the portion of the L1 response that was eliminated by the rectification. Note the faster and stronger response to a dark-to-light transition. **(C)** Simulated responses to motion-blurred light and dark edges by an L2 pathway-like model incorporating L2-type dynamic feedback followed by half-wave rectification. Note the faster and stronger response to light-to-dark transition as well as the faster response to a dark-to-light transition.

**Figure S5.**
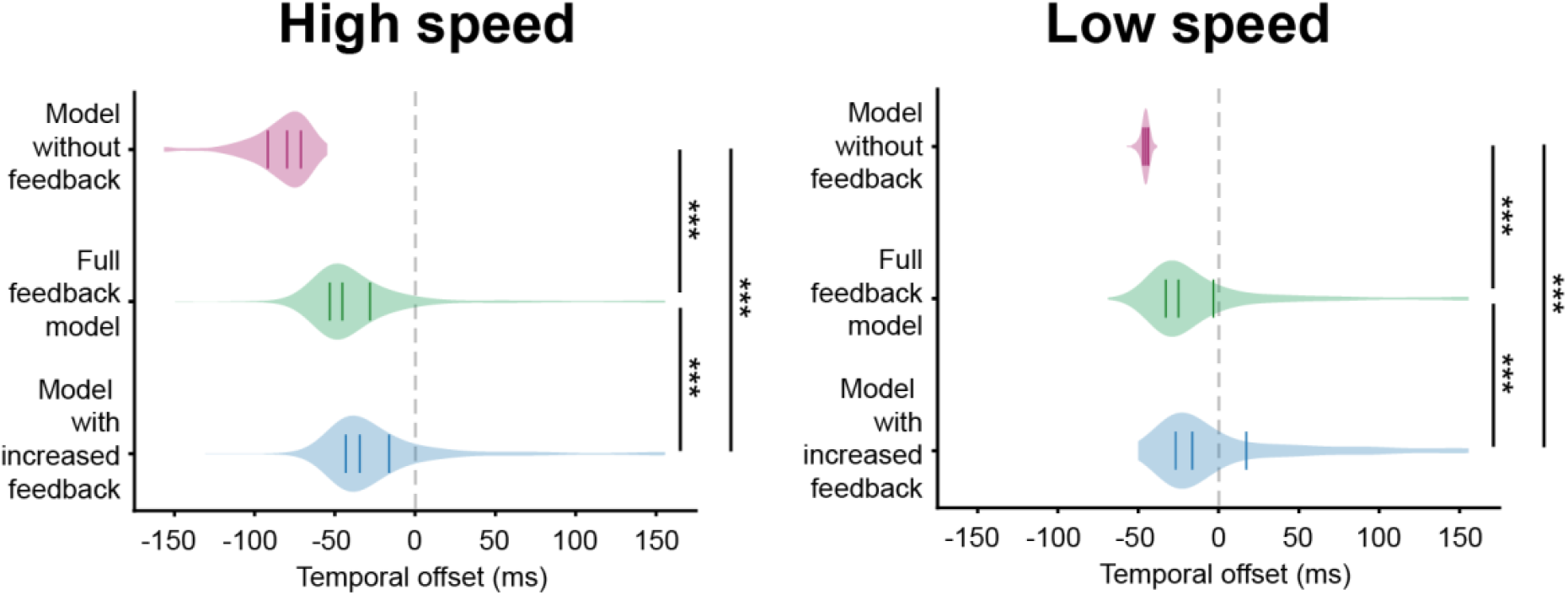
Increased feedback improves deblurring at fast image speeds for horizontal motion. Fig. S5 Image description Violin plots of the temporal offset between the natural image and model output at high (left, 600 °/s) and low (right, 150 °/s) image speeds, using *w* = 0 (model without feedback), *w* = 12.7 (full feedback model), and *w* = 24 (model with increased feedback). Images were blurred with motion in the horizontal direction. Solid lines indicate the lower quartile, median, and upper quartile. Dotted gray line indicates temporal offset = 0 ms. *** p<0.001, Wilcoxon signed-rank test.

**Figure S6.**
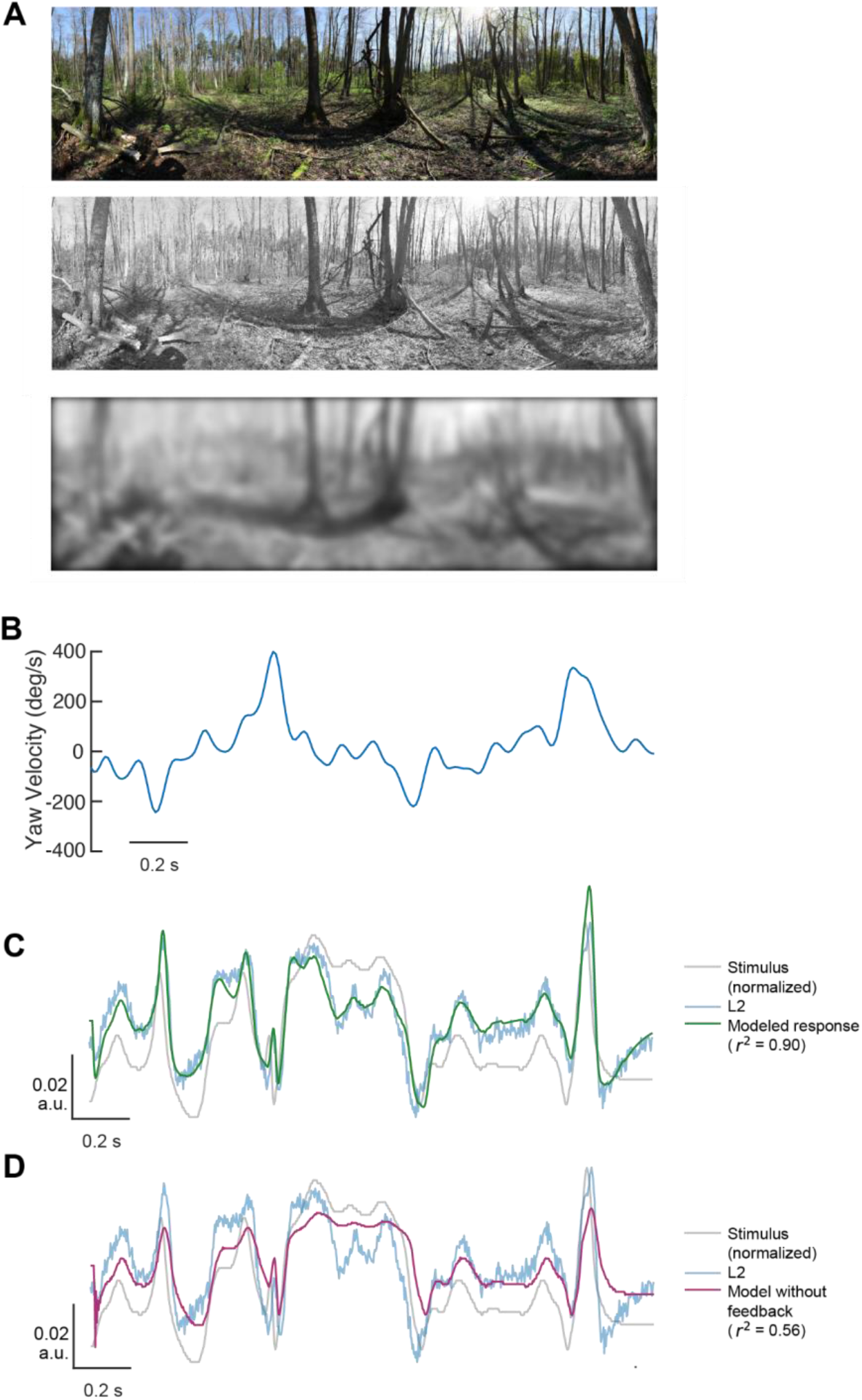
Feedback improves model predictions of L2 responses to a naturalistic stimulus when fit to those responses. Fig. S6 Image Description **(A)** A natural image was converted to grayscale without gamma correction and blurred based on the optics of the fly eye. **(B)** Yaw velocity sequence from a trajectory of a walking fly that was used to determine the sequence of pixels in the natural image from which to draw stimulus intensity values. **(C)** Full model of L2 responses to the naturalistic stimulus using parameters fitted to the naturalistic stimulus response (overlaid on the stimulus inverted across the x-axis) **(D)** Modeled responses without feedback to the naturalistic stimulus using parameters fitted to the naturalistic stimulus response (overlaid on the stimulus inverted across the x-axis)

## STAR Methods

## KEY RESOURCES TABLE

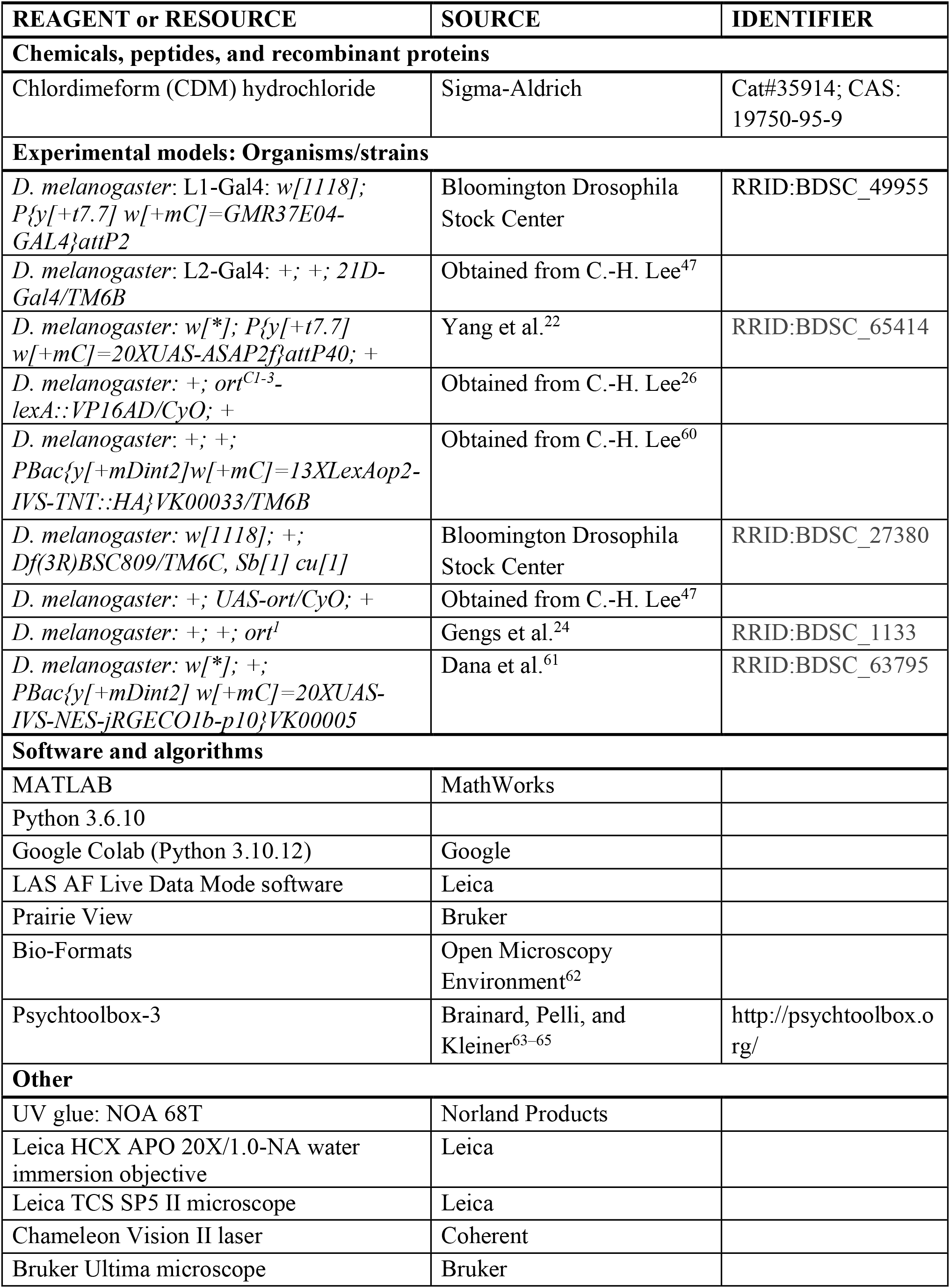
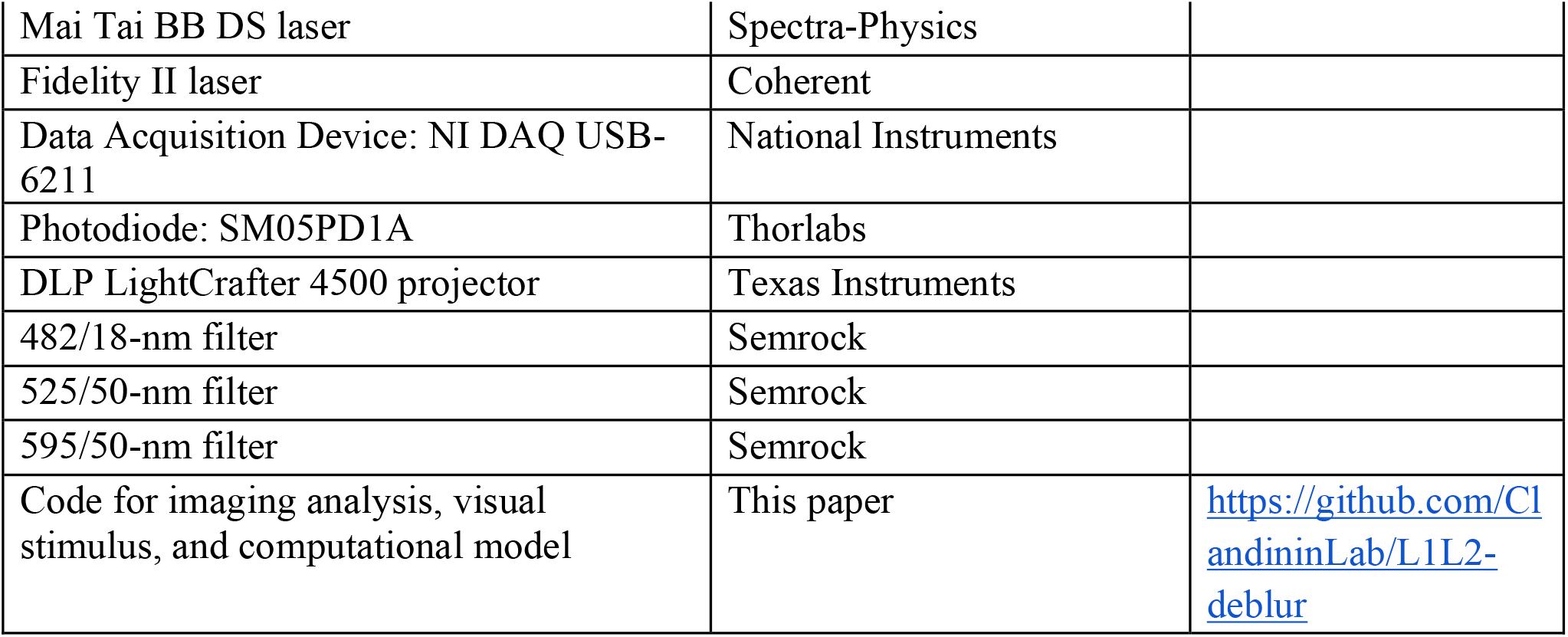

## EXPERIMENTAL MODEL AND STUDY PARTICIPANT DETAILS

Flies were raised on standard molasses food at 25 °C on a 12/12-h light-dark cycle. Female flies of the following genotypes were used:

- L1>>ASAP2f (Figures 1 and 5): *w/+; UAS-ASAP2f/+; GMR37E04-GAL4/+*
- L2>>ASAP2f (Figures 1 and 5): *+; UAS-ASAP2f/+; 21D-GAL4/+*
- *ort*^*C1-3*^*>>TNT* experimental (Figure 2): *+; UAS-ASAP2f/ort*^*C1-3*^*-lexA::VP16AD; 21D-GAL4/13XLexAop2-IVS-TNT::HA*
- *ort*^*C1-3*^*-lexA* control (Figure 2): *+; UAS-ASAP2f/ort*^*C1-3*^*-lexA::VP16AD; 21D-GAL4/+*
- *lexAop-TNT* control (Figure 2): *+; UAS-ASAP2f/+; 21D-GAL4/13XLexAop2-IVS-TNT::HA*
- *Df(3R)BSC809/+, L2>>ort* control (Figure 2): *w/+; UAS-ASAP2f/UAS-ort; 21D-GAL4/Df(3R)BSC809*
- *ort*^*1*^*/+, L2>>ort* control (Figure 2): *+; UAS-ASAP2f/UAS-ort; 21D-GAL4, ort*^*1*^*/+*
- *ort*^*1*^*/Df(3R)BSC809, L2>>ort* experimental (Figure 2): *+; UAS-ASAP2f/UAS-ort; 21D-GAL4, ort*^*1*^*/Df(3R)BSC809*
- *ort*^*1*^*/Df(3R)BSC809* control (Figure S1): *+; UAS-ASAP2f/+; 21D-GAL4, ort*^*1*^*/Df(3R)BSC809*
- L2>>ASAP2f, jRGECO1b (Figure 6): *+; UAS-ASAP2f/+; 21D-GAL4/UAS-jRGECO1b*

## METHOD DETAILS

### Fly preparation

Flies were prepared for imaging as previously described^22,66^. Briefly, flies were collected on CO_2_ 1-2 days post-eclosion and imaged 5-8 days post-eclosion. The flies were cold anesthetized, positioned in a custom-cut hole in a steel shim, and immobilized with UV glue (NOA 68T from Norland Products). The back of the head capsule was dissected with fine forceps to expose the left optic lobe and perfused with a room temperature, carbogen-bubbled saline-sugar solution (modified from ref.^67^). The saline composition was as follows: 103 mM NaCl, 3 mM KCl, 5 mM TES, 1 mM NaH_2_PO_4_, 4 mM MgCl_2_, 1.5 mM CaCl_2_, 10 mM trehalose, 10 mM glucose, 7 mM sucrose, and 26 mM NaHCO_3_. The pH of the saline equilibrated near 7.3 when bubbled with 95% O_2_/5% CO_2_. For the CDM experiments, a 200 mM stock solution was made by dissolving the powder in distilled water and stored at 4 °C. Saline-sugar solution with 20 μM CDM was made fresh from the stock solution daily. Each fly was imaged for up to 1 h.

### Two-photon imaging

The L1 arbor was imaged in medulla layer M1, and the L2 arbor was imaged in medulla layer M2.

In Figures 1-3, and 5, we used a Leica TCS SP5 II system with a Leica HCX APO 20X/1.0-NA water immersion objective and a Chameleon Vision II laser (Coherent). Data were acquired at a frame rate of 82.4 Hz using a frame size of 200×20 pixels, pixel size of 0.2 μm× 0.2 μm, a line scan rate of 1400 Hz, and bidirectional scanning. ASAP2f was excited at 920 nm with 5-15 mW of post-objective power, and photons were collected through a 525/50-nm filter (Semrock). Imaging was synchronized with visual stimulation (see below) using triggering functions in the LAS AF Live Data Mode software (Leica), a Data Acquisition Device (NI DAQ USB-6211, National Instruments), and a photodiode (Thorlabs, SM05PD1A) directly capturing the timing of stimulus frames. Data were acquired on the DAQ at 5000 Hz.

In Figure 6, we used a Bruker Ultima system with a Leica HCX APO 20X/1.0-NA water immersion objective and a Mai Tai BB DS laser (Spectra-Physics). Data were acquired at a frame rate of 354.4 Hz using a frame size of 200×40 pixels, pixel size of 0.2 μm×0.2 μm, 8 kHz line scan rate, and bidirectional resonant scanning. ASAP2f was excited at 920 nm with 5-15 mW of post-objective power, and photons were collected through a 525/50-nm filter (Semrock). To identify cells that respond to a search stimulus during imaging (see below), jRGECO1b signals were excited at 1070 nm using a Fidelity II laser (Coherent) and collected through a 595/50-nm filter (Semrock). Imaging was synchronized with visual stimulation (see below) using a photodiode (Thorlabs, SM05PD1A) connected to the Bruker system to collect the timing of stimulus frames.

### Visual stimulation

Visual stimuli were generated with custom-written software using MATLAB (MathWorks) and Psychtoolbox-3^63–65^ and presented using the blue LED of a DLP LightCrafter 4500 projector (Texas Instruments), as previously described^66^. The projector light was filtered with a 482/18-nm bandpass filter (Semrock) and refreshed at 300 Hz. The stimulus utilized 6 bits/pixel, allowing for 64 distinct luminance values. The radiance at 482 nm was approximately 78 mW·sr^-1^·m^-2^. In Figures 1-3 and 5, the stimulus was rear-projected onto a screen positioned approximately 8 cm anterior to the fly that spanned approximately 70° of the fly’s visual field horizontally and 40° vertically. In Figure 6, the stimulus was rear-projected onto a screen positioned approximately 6 cm anterior to the fly that spanned approximately 80° of the fly’s visual field horizontally and 50° vertically. All stimuli also included a small square simultaneously projected onto a photodiode to synchronize visual stimulus presentation with the imaging data. Stimulus presentation details were saved in MATLAB .mat files to be used in subsequent processing.

The visual stimuli used were:

- Search stimulus: alternating light and dark flashes, each 300 ms in duration, were presented at the center of the screen (8° from each edge of the screen from the perspective of the fly). The responses to this stimulus were used during analysis to exclude cells with receptive fields located at the edges of the screen, which would have uneven center-surround stimulation during full-field stimuli. Cells with receptive fields at the edge of the screen would have opposite-signed responses to the search stimulus and the full-field stimulus, whereas cells with receptive fields at the center of the screen would have responses with the same sign to the two stimuli. The search stimulus was presented for 61 s per field of view for data shown in Figures 1-3 and 5, and 30 s per field of view for data shown in Figure 6.
- Impulse stimulus (Figures 1-3 and 5): 20 ms light and dark flashes were presented over the entire screen with 500 ms gray interleaves between flashes. The light and dark flashes were randomly chosen at each presentation. The Weber contrast of the flashes relative to the gray interleave was 1. This stimulus was presented for 121 s per field of view.
- Naturalistic stimulus (Figure 6): a 2 second sequence of varying intensities extracted from a moving panoramic natural image was presented over the entire screen. The natural image^68^ was processed to remove gamma correction and converted to grayscale. A Gaussian filter was applied to the image to simulate optical blur due to the angular resolution of the fly eye^38,69^. To simulate motion blur, we used a time-varying sequence of yaw rotation angles converted from the angular velocity trajectory of a fly freely walking on a flat surface^70,71^. The intensity of the pixels corresponding to the yaw angle sequence were obtained from the image and normalized to a range of 0 to 1. This stimulus was presented for 360 s per field of view.

### In vivo imaging data analysis

Imaging data were analyzed as previously described^66^. Time series were read into MATLAB using Bio-Formats and motion-corrected by maximizing the cross-correlation in Fourier space of each image with the average of the first 30 images. ROIs (regions of interest) corresponding to individual cells were manually selected from the average image of the whole time series, and the fluorescence of a cell over time was extracted by averaging the pixels within the ROI in each imaging frame. The mean background value was subtracted either frame by frame (Leica) or line by line (Bruker) to correct for visual stimulus bleedthrough. To correct for bleaching, the time series for each ROI was fitted with the sum of two exponentials, and the fitted value at each time t was used as F_0_ in the calculation of ΔF/F = (F(t) – F_0_)/F_0_. For the search stimulus, all imaging frames were used to compute the fit, thereby placing ΔF/F = 0 at the mean response after correction for bleaching. For the impulse stimulus, only imaging frames that fell in the last 25% of the gray period were used to fit the bleaching curve; this places ΔF/F = 0 at the mean baseline the cell returns to after responding to the flash instead of at the mean of the entire trace (which is inaccurate when the responses to the light and dark flashes are not equal and opposite). Time series were manually inspected and discarded if they contained uncorrected movement. For the search and impulse stimuli, the stimulus-locked average response was computed for each ROI by reassigning the timing of each imaging frame to be relative to the stimulus transitions and then averaged across a window of 8.33 ms.

This effectively resampled our data from 82.4 Hz to 120 Hz. For the naturalistic stimulus, responses were recorded at 354.4 Hz so the stimulus-locked average response was simply computed by assigning the first frame after the start of each stimulus to t=0 and averaging across trials.

As the screen on which stimuli were presented did not span the fly’s entire visual field, only a subset of imaged ROIs experienced the stimulus across the entire extent of their spatial receptive fields. These ROIs were identified based on having a response of the appropriate sign to the search stimulus. ROIs lacking a response to this stimulus or having one of the opposite sign were not considered further, except for the *ort*^*1*^*/Df(3R)BSC809* control condition (Figure S1). As L2 did not respond to either the search stimulus or the impulse stimulus in this genotype, all ROIs not discarded for motion artifacts were considered valid. All traces are presented as the mean ± 1 SEM across all of the valid ROIs. Because ASAP2f decreases in brightness upon depolarization, ΔF/F y-axes are in units of negative % ΔF/F so that depolarization is up and hyperpolarization is down.

The metrics quantifying the impulse responses are defined as:

- Phase 1 Area: The area under the curve from the time when the stimulus-evoked response first starts to when the response equals zero between the first and second phases. The start of the response was defined as the first time when the change in the response (ΔF/F, arbitrary units, or normalized amplitude) between one frame and the next is more than the change between that frame and the previous one plus 10%. After the signed area is computed, the sign is flipped if necessary such that the expected direction of the response is defined as positive (e.g. L1 and L2 initially depolarize to dark, which is a negative ΔF/F, so the phase 1 area for the dark flash is multiplied by −1).
- Phase 2 Area: The area under the curve from the time when the response equals zero between the first and second phases to 0.25 seconds after the start of the stimulus. As with the Phase 1 Area, positive is defined as the area in the expected direction of the response.
- Area Phase 2/Phase 1: The Phase 2 Area divided by Phase 1 Area.
- Phase 1 t_peak_: The time when the stimulus-evoked response first starts to the time when magnitude of the first phase is maximal.
- Phase 2 t_peak_: The time when the stimulus-evoked response first starts to the time when the magnitude of the second phase is maximal, unless the Phase 2 Area was not statistically significantly different from 0, in which case it was not reported.

The responses of all of the valid ROIs of a particular genotype or condition were bootstrapped (sampled with replacement with the sample size equal to the original sample size) and the quantification metrics were computed on the mean trace; this was repeated 10,000 times to calculate the 95%, 99%, and 99.9% confidence intervals. The mean and 95% confidence interval was plotted. The confidence intervals were used to test for a statistically significant difference between two genotypes or conditions: if the confidence intervals did not overlap, then the difference was statistically significant at the corresponding α level (e.g. 95% for α = 0.05, 99% for α = 0.01, and 99.9% for α = 0.001). To correct for multiple comparisons (Figure 2), Bonferroni correction was applied, and the confidence intervals corresponding to particular α levels were adjusted accordingly.

### Computational model

To maximize interpretability, a feedback model consisting of two differential equations was used: one corresponding to an effective voltage variable that captures the membrane potential of L1 or L2 (Eq. 1) and the second to the dynamics of the recurrent feedback circuit (Eq. 2A). The simplified linear form of the model is:

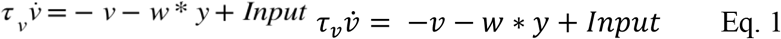

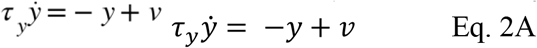

where τ_*v*_ and τ_*y*_ describe time constants for *v* and *y*, the neural response and the feedback element, respectively, and *w* describes the strength of the feedback. However, this linear form of our model is not capable of capturing the strong asymmetry in the impulse responses to light and dark, in turn reflecting differences in the ability of light and dark to drive feedback. Therefore, Eq. 2A was modified to capture this asymmetry (Eq. 2B):

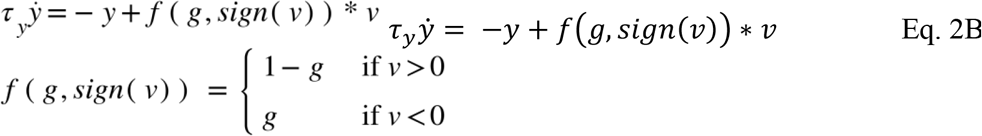

Accordingly, when *g* is equal to one, only light drives the feedback element; when *g* is equal to zero, only dark responses drive the feedback element, and when *g* = 0.5 light and dark responses drive feedback equally. In addition, to account for the asymmetry of the first phase of response in which the feedback plays little role, we assume that the input term in Eq. 1 is weighted differently for light and dark responses as follows:

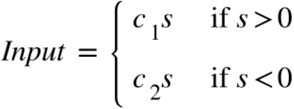

Where *s* denotes the stimuli and *c*_*1*_, *c*_*2*_ are constant weights. Additionally, an affine transformation is performed on the output to match the experiment scale. All parameters associated with the computational model were fitted through gradient descent to minimize the mean square error (MSE) between the model’s predictions and the experimental responses to the light and dark flashes. The model was simulated using the Euler method to match the experimental sampling frequency. The Adam optimizer was used with an initial learning rate of 0.005 for the first 3000 steps, followed by a reduced learning rate of 0.001 for an additional 2000 steps.

When fitting the data, a latency difference between the model output and the experimental data was found. To reconcile this latency difference, the inputs were delayed by *d* milliseconds. A range of delays were swept, and *d* = 16.7 ms explained the experimental data the best. This delay approximates the delays associated with phototransduction, which were not accounted for in the model^51^.

In Figure 3B, the fitted values for L1 were: *τ*_*v*_= 0.0127 s, *τ*_*y*_= 0.165 s, *w*=8.00, *g* = 0.0277. The values used for L2 were: *τ*_*v*_= 0.0131 seconds, *τ*_*y*_= 0.561 s, *w* = 12.7, *g* = 0.169. Figure 3C used L1 and L2 parameter values, aside from *w*, which was systematically swept across a range of values. The models in Figure 4 used the values corresponding to L2. Qualitatively similar results held when using parameters corresponding to L1. In Figure 5, the increased feedback model had *w* = 10.40 for L1 and *w* = 23.63 for L2, obtained by fitting the model to the CDM responses. All other model parameters were identical to those used in Figure 3. In Figures 3C, S3, and S4, the model was simulated with an input of length 0.47 seconds starting at 0.155 seconds. In Figure S4, the model of the L1 pathway was implemented by adding a subsequent sign-inversion followed by half-wave rectification; the model of the L2 pathway was implemented by adding a subsequent half-wave rectification.

In Figures 6C and 6D, the L2 parameters fitted from the impulse responses were used, and only the final affine transformation was refit. The model without feedback was obtained by setting *w* = 0. The full model was also fit to the naturalistic stimulus response, as shown in Figure S6. The model parameters were initialized to be the same as the L2 parameters, and then the Adam optimizer was used to train the model with an initial learning rate of 0.002 for 2000 steps, followed by a reduced learning rate of 0.001 for an additional 1000 steps. The fitted parameters were: *τ*_*v*_= 0.0043 seconds, *τ*_*y*_= 0.417 seconds, *w* = 12.5, *g* = 0.275. The model without feedback was trained with *w* = 0, and the initial learning rate and the reduced learning rate were changed to 0.0002 and 0.0001 respectively. The fitted parameters for the model without feedback were: *τ*_*v*_= 0.00176.

For the convolutional model, the light or dark impulse responses of L2 were used depending on the input sign:

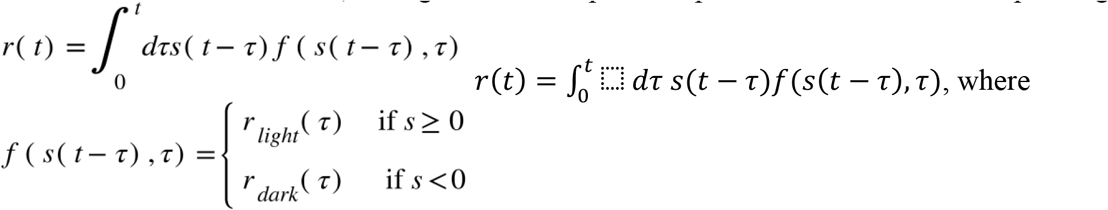

and *r*_*light/dark*_ (*τ*) are L2 responses to the light or dark impulse stimulus.

The van Hateren dataset of natural images was used to conduct simulations^37^. To preprocess and subsample the dataset, we first standardized (Z-scored) each image, and then analyzed the standardized images before and after binarizing them using a threshold of 0. Contrast values were determined for each standardized and thresholded image by taking the standard deviation of each column (vertical) or row (horizontal) and averaging those values. These contrast values were then used to subsample the highest-contrast images in the dataset such that the standardized and thresholded images both had contrast values in the top quartile. Our simulation results only hold for high-contrast images. Translating these natural images to input was done by incorporating photoreceptor optics and rigid translational motion, as in ref.^38^. In brief, this transformation comprised two steps: a spatial blur introduced by the optics of the fly visual system, modeled by a Gaussian blur, and a temporal blur corresponding to the visual scene moving during the non-zero photoreceptor integration time, modeled by a temporal exponential blur. An image speed of 300 degrees per second was used for Figure 4. 150 and 600 degrees per second were used for low and high speeds, respectively, in Figure 5. The output image was constructed by first translating the original image into a spatially and temporally blurred image. Then, each column (vertical) or row (horizontal) of this transformed image was used as the temporal input to the feedback model. A final output image was constructed by combining the outputs of the model running on each column or row. Denote the origin image as *X* and the model reformed image as *Y*. We infer the latency between the origin image and the model reconstructed image from the following optimization for vertical moving direction:

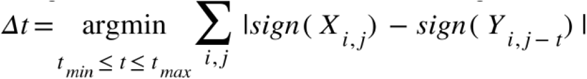

where we choose *t*_*min*_ = −156 ms and *t*_*max*_ = 156 ms. For horizontal moving direction, we use the transposed image in the above equation. To reduce the boundary effect, we only sum *j* from 400 to 800 pixels (for vertical ones) and 400 to 1336 pixels (for horizontal ones). To save compute time, we sampled *i* from 200 to 400 with a stride of 2.

### Idealized impulse responses (Figure S2)

Idealized impulse responses were constructed in MATLAB by defining the time at which the response starts relative to the “start of the stimulus” as well as the time at which the response reaches peak amplitude for both the first and second phases and their amplitudes at those times. Linear interpolation defined the values between the response start and the peak of the first phase as well as between the peaks of the two phases. The decay of the second phase was a single exponential decay with the same decay constant across all of the idealized impulse responses. The peak amplitude of the first phase was held constant, and the peak amplitude of the second phase was varied to change the relative size of the second phase. The monophasic impulse response had a second phase peak amplitude of 0. The parameters were chosen to approximate the measured impulse responses and produce the desired area ratio but were not directly fit. The sample rate was 120 Hz, matching that of the measured responses. The magnitude across frequencies was computed by calculating the absolute value of the Fourier transform of idealized impulse responses normalized to energy. The simulated response to a 250 ms dark step off of gray was computed by convolving the impulse responses with that stimulus.

## QUANTIFICATION AND STATISTICAL ANALYSIS

All statistical details are described briefly in the figure legends and in depth in the Method Details. Statistical significance was defined at a = 0.05, and Bonferroni correction for multiple comparisons was applied when appropriate. Statistical analyses were performed in MATLAB and Python.

## RESOURCE AVAILABILITY

### Lead contact

Further information and requests for resources and reagents should be directed to and will be fulfilled by the lead contact, Helen Yang.

### Materials availability

This study did not generate new unique reagents.

### Data and code availability

- All data reported in this paper will be shared by the lead contact upon request.
- All original code has been deposited at Github and listed in the key resources table.
- Any additional information required to reanalyze the data reported in this paper is available from the lead contact upon request.

## Appendix Image Descriptions

**Fig. 1 Long Description**

A. Two schematized line plots connected by an inhibitory arrow going from R1-6 to L1/L2
  - Top (R1-6): A trace begins at the x-axis, sharply increases above the axis to a large maximum peak, and then gradually decreases back down to the x-axis.
  - Bottom (L1/L2): A trace begins at the x-axis, sharply decreases below the axis to a large minimum trough, sharply increases to a small maximum peak above the x-axis, and then gradually decreases back down to the x-axis.
B. Anatomy diagram with regions labeled lamina and medulla. The more distal lamina region includes the axonal arbors of R1-6 adjacent to the dendritic arbors of L1/L2. The more proximal medulla region includes the axonal arbors of L1 and L2. L1 has two axonal arbors (the more proximal one of which is boxed), and L2 has one axonal arbor (which is also boxed).
C. Schematized line plot exemplifying how the stimulus-aligned voltage responses will be displayed in panels D-E and quantified in panel F. The y-axis is −ΔF/F (in %); it is inverted so that depolarization is up and hyperpolarization is down. The x-axis is time (in seconds). A single trace begins at a baseline “0%” ΔF/F level, goes downward to a sharp trough (1st peak), then upward to a shallower peak above baseline (2nd peak), and then gradually returns to baseline. The area between the trace and the baseline axis is labeled Phase 1 Area and Phase 2 Area, corresponding to the two “lobes” formed by the 1st and 2nd peaks, respectively. The lengths of time it takes to reach the two peaks are labeled Phase 1 t_peak_ and Phase 2 t_peak_. Above this trace, there is a thin rectangular box representing the visual stimulus, with a small segment at the beginning shaded white indicating the short impulse flash, and a longer segment shaded gray indicating the mean gray interleaf period. The white segment spans approximately half the length of time it takes to reach Phase 1 t_peak_. (For clarity, peaks that are oriented downward in plots will be referred to as “troughs” or “minima” in image descriptions. Peaks that are oriented upward will be referred to as “peaks” or “maxima.” Note that this is the opposite of the numeric minima and maxima, as negative ΔF/F values are oriented upward and positive ΔF/F values are oriented downward.)
D. Two line plots showing the average responses of L1 to light and dark flashes (n = 103 cells, from 13 flies). For each plot, the stimulus schematic is displayed at the top, and the response trace is underneath, as in C. The x-axis is in time, spanning 0 to 0.5 second. The y-axis is in −ΔF/F, spanning +8% to −4% (values provided in this description are approximate).
  - The light response begins at baseline (0%), goes downward to a sharp trough at +6%, and then goes upward back to the baseline, where it remains for the rest of the gray period of the stimulus. The width of the first trough is about twice the duration of the light flash.
  - The dark response begins at baseline (0%), goes upward to a sharp peak at −3%, goes downward past baseline to a trough at +3%, and then gradually goes upward back to baseline, reaching 0% near the end of the gray period. The width of the first peak is about twice the duration of the light flash.
E. As in D, but for L2 (n = 68 cells, from 14 flies).
  - The light response begins at baseline (0%), goes downward to a sharp trough at +5%, goes upward past baseline to a small peak at −1%, and then gradually goes downward back to baseline, reaching the baseline around 0.15 second. The width of the first trough is about twice the duration of the light flash.
  - The dark response begins at baseline (0%), goes upward to a sharp peak at −3%, goes downward past baseline to a trough at +2%, and then gradually goes upward back to baseline, reaching baseline near the end of the gray period. The width of the first peak is about twice the duration of the light flash.
F. Five panels of dot-and-whisker plots quantifying the impulse responses shown in D-E. Each panel has four sets of dot-and-whiskers corresponding to the mean and 95% confidence interval from L1 light, L1 dark, L2 light, and L2 dark. Approximate dot and whisker values (and their statistical comparisons) are summarized in the following table as “dot ± whiskers” or “dot ± (−lower, +upper)” when whiskers are asymmetric:

| | Phase 1 Area (-<br>%$\Delta F/F$ * s) | Phase 2 Area (-<br>%$\Delta F/F$ * s) | Area Phase 2<br>/ Phase 1 | Phase 1 $t_{peak}$ (s) | Phase 2 $t_{peak}$ (s) |
| --- | --- | --- | --- | --- | --- |
| **L1 light** | 0.13 ± 0.015 | -0.005 ± 0.01 | 0 ± negligible | 0.017 ± (-0.008,<br>+negligible) | No data |
| **L1 dark** | 0.045 ± 0.01 | 0.23 ± 0.03 | 5.2 ± 0.9 | 0.016 ± (-0.007,<br>+0.01) | 0.04 ± (-0.005,<br>+0.01) |
| **L2 light** | 0.1 ± 0.005 | 0.05 ± 0.01 | 0.050 ±<br>negligible | 0.017 ± (-0.008,<br>+negligible) | 0.04 ± (-0.005,<br>+0.01) |
| **L2 dark** | 0.050 ± 0.005 | 0.12 ± 0.015 | 2.4 ± 0.4 | 0.016 ± (-0.007,<br>+0.01) | 0.05 ± (-0.01,<br>+0.02) |
| **\*\*\***<br>**(p<0.001)** | L1 light & dark;<br>L2 light & dark;<br>L1 & L2 light | All pairs | All pairs | None | None |
| **n.s.**<br>**(p>0.05)** | L1 & L2 dark | None | None | All pairs | L2 light & dark<br>L1 & L2 dark |

<u>Back to Fig. 1</u>

**Fig. 2 Long Description**

Title above A-C: “L2 responses to photoreceptor input only (ort-neuron silencing)”

Title above D-F: “L2 responses to photoreceptor input and recurrent feedback (L2 ort rescue)”

A. Five circles representing photoreceptors, L2 cells, and three types of feedback neurons. There are arrows representing the following possible circuit pathways, as well as X’s over some arrows representing silenced pathways under the ort-neuron silencing condition (ort driver expressing TNT):
  - Photoreceptors to L2
  - Photoreceptors to lateral feedback neurons
  - Lateral feedback neurons to L2 (X, silenced)
  - Lateral feedback neurons to downstream feedback neurons (X, silenced), then to L2
  - L2 to recurrent feedback neurons (X, silenced), then back to L2
B. Two panels of line plots showing the average responses of L2 to light and dark flashes under the ort-neuron silencing condition. The x-axis is in time, spanning 0 to 0.5 second. The y-axis is in −ΔF/F, spanning +8% to −4% (values provided in this description are approximate).
  - Each panel has three overlaid traces for the following genotypes:
    - ort-lexA control (n = 129 cells from 11 flies)
    - lexAop-TNT control (n = 96 cells from 9 flies)
    - ort-lexA, lexAop-TNT experimental (n = 116 cells from 9 flies)
  - Light impulse panel:
    - The two control traces are largely the same; both begin at 0%, go downward to a sharp trough at +5%, go upward past 0 to a small peak at −1%, and then gradually go downward back to 0, reaching 0 around 0.2 second. The width of the troughs is around twice the duration of the light flash.
    - The experimental trace begins at 0%, goes downward to a sharp trough at +6%, gradually goes upward back to 0 (without crossing 0), reaching 0 at around 0.2 second. The width of the trough is around four times the duration of the light flash.
  - Dark impulse panel: All three traces are initially largely the same; all begin at 0%, go upward to a sharp peak at −2.5%, and go downward past 0 to a trough at +2%. The control traces immediately start returning to 0 gradually, reaching 0 at around 0.4 second. In contrast, the experimental trace perdures at +2% for around 0.2 second before gradually returning to 0, reaching 0 near the end of the gray period.
C. Five panels of dot-and-whisker plots quantifying the impulse responses shown in B. Each panel has dot-and-whiskers corresponding to the means and 95% confidence intervals from six conditions, which include light and dark responses from three genotypes: ort-lexA control (Control 1), lexAop-TNT control (Control 2), and ort-lexA, lexAop-TNT (experimental). Approximate dot and whisker values (and their statistical comparisons) are summarized in the following table as “dot ± whiskers” or “dot ± (−lower, +upper)” when whiskers are asymmetric:

D. Five circles representing photoreceptors, L2 cells, and three types of feedback neurons. There are the same arrows as in A representing possible circuit pathways, but without X’s, as there is no presynaptic silencing under the L2 ort rescue condition (ort null background with L2-Gal4 > UAS-ort). The L2 circle has membrane receptors representing the presence of ort receptors. The lateral feedback neuron circle has membrane receptors that are crossed out, representing the absence of ort receptors in these cells. Thus, the pathway from photoreceptors to L2 is intact, while the pathway from photoreceptors to lateral feedback neurons is blocked.
E. As in B, but for the L2 ort rescue condition:
  - Each panel has three overlaid traces for the following genotypes: (all of which additionally include L2-Gal4, UAS-ort):
    - Df(3R)BSC809/+ control (n = 94 cells from 7 flies)
    - ort^1^/+ control (n = 59 cells from 5 flies)
    - Df(3R)BSC809/ort^1^ experimental (n = 58 cells from 6 flies)
  - Light impulse panel: All three traces are largely the same; all begin at 0%, go downward to a sharp trough at around +5%, go upward past 0 to a small peak at −1%, and then gradually go downward back to 0, reaching 0 around 0.15 second. The width of the first trough is around twice the duration of the light flash.
  - Dark impulse panel: All three traces are initially largely the same; all begin at 0% and go upward to a sharp peak at −3%. The control traces go downward past 0 to a trough at +2%, whereas the experimental trace only goes to +1%. All three traces then gradually return to 0, reaching 0 at about 0.3 second. The width of the first peak is around twice the duration of the light flash.
F. As in C, but quantifying the L2 ort rescue impulse responses shown in E. Each panel includes light and dark responses from three genotypes (all of which additionally include L2-Gal4, UAS-ort): Df(3R)BSC809/+ control (Control 1), ort^1^/+ control (Control 2), and Df(3R)BSC809/ort^1^ (experimental).

| | **Phase 1 Area**<br>(-%$\Delta F/F$ * s) | **Phase 2 Area**<br>(-%$\Delta F/F$ * s) | **Area Phase 2 / Phase 1** | **Phase 1 $t_{peak}$ (s)** | **Phase 2 $t_{peak}$ (s)** |
| --- | --- | --- | --- | --- | --- |
| **Control 1, light** | 0.11 ± 0.01 | 0.05 ± 0.02 | 0.5 ± negligible | 0.016 ± (-0.008, +negligible) | 0.08 ± 0.025 |
| **Control 2, light** | 0.09 ± 0.01 | 0.06 ± 0.02 | 0.8 ± negligible | 0.016 ± (-0.008, +negligible) | 0.06 ± (-0.01, +0.02) |
| **Experimental, light** | 0.25 ± 0.03 | 0.005 ± negligible | 0 ± negligible | 0.025 ± negligible | No data |
| **Control 1, dark** | 0.050 ± negligible | 0.13 ± 0.03 | 3.2 ± 1.5 | 0.024 ± (-0.008, +negligible) | 0.04 ± negligible |
| **Control 2, dark** | 0.055 ± negligible | 0.13 ± 0.02 | 3.0 ± 1 | 0.016 ± (-negligible, +0.008) | 0.04 ± (-negligible, +0.02) |
| **Experimental, dark** | 0.055 ± negligible | 0.20 ± 0.03 | 4.4 ± 1.5 | 0.025 ± negligible | 0.1 ± (-0.02, +0.08) |
| **\*\*\* (p<0.001)** | Light responses | Light responses | Light responses | Light responses | None |
| **\*\* (p<0.01)** | None | Dark responses | None | None | Dark responses |
| **n.s. (p>0.05)** | Dark responses | None | Dark responses | Dark responses | None |

|  | Phase 1 Area<br>(-%ΔF/F * s) | Phase 2 Area<br>(-%ΔF/F * s) | Area Phase 2 /<br>Phase 1 | Phase 1 t<sub>peak</sub> (s) | Phase 2 t<sub>peak</sub><br>(s) |
| --- | --- | --- | --- | --- | --- |
| **Control 1,**<br>**light** | 0.11 ± 0.01 | 0.04 ± 0.01 | 0.4 ± negligible | 0.017 ±<br>negligible | 0.09 ± (-0.05,<br>+0.4) |
| **Control 2,**<br>**light** | 0.10 ± 0.01 | 0.06 ± 0.02 | 0.6 ± 0.1 | 0.011 ± (-0.003,<br>+0.007) | 0.05 ±<br>negligible |
| **Experimental,**<br>**light** | 0.09 ± 0.01 | 0.04 ± 0.02 | 0.5 ± 0.1 | 0.017 ±<br>negligible | 0.05 ±<br>negligible |
| **Control 1,**<br>**dark** | 0.050 ± 0.005 | 0.14 ± 0.02 | 3.0 ± 0.5 | 0.017 ±<br>negligible | 0.04 ±<br>negligible |
| **Control 2,**<br>**dark** | 0.055 ± 0.01 | 0.14 ± 0.02 | 2.5 ± 0.5 | 0.016 ± (-0.008,<br>+negligible) | 0.04 ±<br>negligible |
| **Experimental,**<br>**dark** | 0.060 ± 0.005 | 0.09 ± 0.02 | 1.5 ± 0.3 | 0.017 ±<br>negligible | 0.04 ±<br>negligible |
| **\* (p<0.05)** | None | Dark responses | Dark responses | None | None |
| **n.s. (p>0.05)** | Light and dark<br>responses | Light responses | Light responses | Light and dark<br>responses | Light and dark<br>responses |

<u>Back to Fig. 2</u>

**Fig. 3 Long Description**

A. Schematic and equations describing the recurrent feedback model:
  - Schematic: Circles and arrows representing the neural circuit simulated by this computational model. The three main elements are the visual input, L1/L2 (*v*), and the feedback element (*y*). L1/L2 and the feedback element have an additional variable describing their temporal properties (*τ*_*v*_ and *τ*_*y*_, respectively) The arrows indicate the following circuit pathways, whose strength is described by additional variables:
    ∘ Visual input feeds forward to L1/L2, which feeds forward to the rest of the visual system
    ∘ L1/L2 also feeds forward to the feedback element with variable strength dependent on *g*
    ∘ The feedback element provides oppositely-signed feedback to L1/L2, with strength *w*
  - Equations:
    - 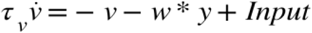
    - 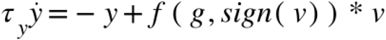
    - 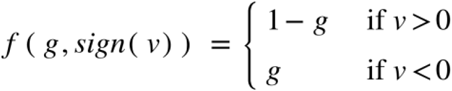
    - *τ*_*y*_ > > *τ*_*ν*_
B. Four panels of line plots showing modeled L1 and L2 responses to light and dark flashes (overlaid on top of the measured impulse responses from Fig. 1D). All four sets of modeled responses overlap closely with the measured responses (except for the measured responses having more noise). There are two subtle differences in all the cases: in the modeled responses, the first peak is slightly rounder, and the second peak has an onset that is less sharp (resulting in a peak that is slightly later and smaller on the onset side).
C. Two panels of line plots, each with eight lines, showing modeled L1 and L2 responses to motion blurred edges with varying feedback strengths (*w* = 0, 1, 2, 4, 8, and 12). The L1 and L2 conditions are qualitatively similar and will be described together, with noticeable differences specified. The y-axis is Activation (A.U.) spanning −1 to 1, and the x-axis is time (s) spanning 0 to 0.8 second. Values provided are approximate.
  - The input is a dotted line that starts at 0, drops down to −1 at 0.15s, and then jumps up to 0.5 at 0.62s.
  - The blurred input is a dotted line that matches the input, except the two transitions (from 0 to −1, and from −1 to 0.5) are slower and smoother, taking 0.1s to transition rather than being instantaneous.
  - The *w* = 0 trace is shaped similarly to the blurred input trace, but is shifted later. Like the input, it also travels from 0 to −1 to 0.5 A.U. The transition from 0 to −1 occurs at 0.18-028s, and the transition from −1 to 0.5 occurs at 0.65-0.75s.
  - For L1, the w = 12 trace initially follows the w = 0 trace, but instead of going all the way down to −1, it stops at −0.25 and turns around, reaching −0.1 at around 0.25s. It stays at −0.1 until 0.65s, and then increases to a peak of 0.9 at 0.75s.
  - For L2, the w = 12 trace is qualitatively similar but initially goes a bit further before stopping at −0.5, and takes a bit longer to get back to −0.1 at around 0.35s.
  - The remaining *w* values have traces that span the range between the *w* = 0 and *w* = 12 traces.
  - An arrow above both plots at around 0.65 emphasizes the difference between the traces at that time point. The larger the *w* value, the higher the activation value is at that time, showing that an increase in *w* results in a more rapid readout of the −1 to 0.5 transition due to the feedback engaged during the initial 0 to −1 transition.

<u>Back to Fig. 3</u>

**Fig. 4 Long Description**

A. A series of eight example images depicting different stages of the computational modeling process:
  i. Natural image (example photo of a skyline from the van Hateren dataset, cropped and in grayscale; the remaining images are all processed versions of this photo)
  ii. Optically blurred image.
  iii. Optically and motion blurred image.
  iv. Thresholded natural image.
  v. Thresholded output of the model without feedback.
  vi. Difference between iv and v (the thresholded natural image and the thresholded output of the model without feedback).
  vii. Thresholded output of the full model with feedback.
  viii. Difference between iv and vi (the thresholded natural image and the thresholded output of the full model with feedback). Note that when comparing the thresholded natural image with the two thresholded model outputs, the full model is more closely aligned with the original image than is the model without feedback.
B. Panel titled Vertical Motion, with two violin plots displaying the temporal offsets generated from the model without feedback and the full model with recurrent feedback. The x-axes are of temporal offset in milliseconds, ranging from −160 to 160. The model without feedback has quartiles at −59, −54, and −51, and a range from −98 to −43 (rounded to the nearest digit). The full feedback model has quartiles at −30, −5, and 47, and a range from −119 to 156. These distributions are significantly different (***, p<0.001).
C. As in B, but titled Horizontal Motion. The model without feedback has quartiles at −63, −59, and −56, and a range from −94 to −44. The full feedback model has quartiles at −42, −34, and −13, and a range from −79 to 156. These distributions are significantly different (***, p<0.001).

<u>Back to Fig. 4</u>

**Fig. 5 Long Description**

A. Two panels labeled High Speed and Low Speed, each with three violin plots displaying the distribution of temporal offsets generated from the model without feedback, the full feedback model, and the model with increased feedback. The x-axes are of temporal offset in milliseconds, ranging from −160 to 160. The following describes the violin plots with statistics rounded to the nearest digit:
  - High speed:
    ∘ The model without feedback has quartiles at −85, −74, and −66, and a range from −156 to −48.
    ∘ The full feedback has quartiles at −39, −16, and 19, and a range from −156 to 156.
    ∘ The model with increased speed has quartiles at −31, −7, and 40, and a range from −121 to 156.
    ∘ All three distributions are significantly different from each other (***, p<0.001).
  - Low speed:
    ∘ The model without feedback has quartiles at −45, −42, and −41, and a range from −64 to −36.
    ∘ The full feedback has quartiles at −21, 5, and 58, and a range from −110 to 156.
    ∘ The model with increased speed has quartiles at −12, 18, and 76, and a range from −106 to 156.
    ∘ All three distributions are significantly different from each other (***, p<0.001).
B. Four panels of line plots showing modeled L1 and L2 responses to light and dark flashes. Each panel has a trace labeled “increased feedback” overlaid on a control trace (which is from Fig. 2B). In all four panels, the control and increased feedback traces overlap closely during the first phase (the initial peak or trough). During the second phase, there is no difference in the L1 light response, but the rest of the responses have larger peak amplitudes with increased feedback. The peak amplitudes are increased by around 25%, 100%, and 50% for L1 dark, L2 light, and L2 dark, respectively.
C. Four panels of line plots showing experimental L1 and L2 responses to light and dark flashes. Each panel has a trace labeled “CDM” overlaid on a control trace (which is from Fig. 1D-E). The x-axis is in time, spanning 0 to 0.5 second. The y-axis is in −ΔF/F, spanning +8% to −4% (values provided in this description are approximate).
  - L1 light response: The CDM trace (n = 56 cells, 13 flies) has a trough at +4%, compared to +6% for the control. The CDM also takes longer to return to 0%, around 0.1s instead of 0.05s for the control.
  - L1 dark response: The CDM trace initially peaks at −2%, compared to −3% for the control, and the trough during the second phase is slightly larger, about 3.5% instead of 3% for the control.
  - L2 light response: The CDM trace (n = 102 cells, 7 flies) has a similar first phase compared to the control, but it has a reduced peak in the second phase, around −0.5% instead of −1% for the control.
  - L2 dark response: The CDM trace has a similar first phase compared to the control, but it has a much larger trough in the second phase, which is around +5% compared to +2% for the control.

<u>Back to Fig. 5</u>

**Fig. 6 Long Description**

A. Bottom: Line plot in which the y-axis is Intensity ranging from 0 to 1, and the x-axis is time ranging from 0 to 2 seconds. A single trace fluctuates smoothly throughout most of the range of light intensities over time, starting mostly at the brighter end of the range until around 0.6s, then traveling to the dimmer end of the range until around 1.1s, then returning to the brighter end of the range until the end, except for a rapid excursion to the dimmer end around 1.7s. The initial bright period has two prominent local minima and two prominent local maxima, whereas the middle dim period and the final bright period have smaller fluctuations, except for the final dim excursion. Note that panel B will specifically highlight three short segments of this stimulus: 1. The steep transition from the initial bright period to the middle dim period, 2. A portion of the middle dim period with several small fluctuations, and 3. The rapid dim excursion toward the end of the stimulus (specifically at the steep dim-to bright transition). Top: Rectangle with variable shading in grayscale visualizing the change in brightness of this naturalistic stimulus over time (as described in the line plot).
B. Top: Line plot with two overlaid traces and scale bars for the x- and y-axes. The x-axis is in time, spanning 2 seconds. The y-axis is in −ΔF/F (inverted such that depolarization is up and hyperpolarization is down), spanning approximately 6%. The first trace is an inverted, normalized version of the naturalistic stimulus (the inversion allows dimming of the stimulus and depolarization of L2 to follow the same direction, and vice versa). The second trace shows the average voltage response of L2 (n = 52 cells, from 7 flies), which generally follows the trajectory of the stimulus, lagging slightly behind in time. Bottom: Three notable segments of the stimulus and response are displayed as insets:
  - In response to a rapid bright-to-dim transition, the L2 response begins to increase (depolarize) after the start of the stimulus transition, but this depolarizing excursion then peaks sooner than the end of the stimulus transition.
  - In response to a period of subtle changes in brightness, L2 responds to each increase or decrease in brightness with relatively large amplitudes.
  - In response to a rapid bright-to-dim stimulus transition, the L2 response begins to decrease (hyperpolarize) after the start of the stimulus transition, but this hyperpolarizing excursion then peaks sooner than the end of the stimulus transition.
C. Line plot with three overlaid traces and scale bars for the x- and y-axes. The y-axis is in arbitrary units, spanning approximately 0.08 a.u., and the x-axis is time, spanning 2 seconds. The first trace is an inverted, normalized version of the naturalistic stimulus. The second trace is a normalized version of the measured response of L2. The third trace is the output of the full recurrent feedback model using the same parameters as for the impulse responses. The measured L2 response and the model output are well correlated (r^2^ = 0.70), and the two traces both generally follow the trajectory of the stimulus, lagging slightly behind in time.
D. Same as C, except the third trace is the output of the model without feedback. This trace is less closely correlated with the measured L2 response (r^2^ = 0.39) than in C, and at times it lags noticeably behind not only the stimulus but also the measured L2 response.
E. Same as C, except the third trace is the output of the convolutional model using the impulse response as a linear filter. This trace is less closely correlated with the measured L2 response (r^2^ = 0.39) than in C, and at times it lags noticeably behind not only the stimulus but also the measured L2 response. Compared to the model without feedback in D, the convolutional model has only subtle differences, following some changes in brightness more rapidly and accurately than in D, but sometimes exaggerating the amplitude of smaller changes relative to those of larger changes.

<u>Back to Fig. 6</u>

**Figure S1 Long Description**

Two line plots showing the average responses of L2 to light and dark flashes in a Df(3R)BSC809/ort^1^ control background (n = 121 cells, from 6 flies). The y-axis is in −ΔF/F, spanning +8% to −4%. The x-axis is in time, spanning 0 to 0.5 second. In both the light and dark flash plots, the control trace remains at baseline (0%) throughout the entire duration of the stimulus.

<u>Back to Fig. S1</u>

**Figure S2 Long Description**

A. Three line plots with x-axes spanning 0 to 0.5s, and y-axes spanning around −1 to 1 A.U., schematizing impulse responses with the following waveforms:
  - Monophasic, Area Ratio = 0: The line goes from 0 to −1, and then back to 0.
  - Biphasic, Area Ratio ∼ 1: The line goes from 0 to −1, crosses 0 to a second peak at 0.2, and then gradually returns to 0.
  - Biphasic, Area Ratio ∼ 2: The line goes from 0 to −1, crosses 0 to a second peak at 0.4, and then gradually returns to 0.
B. Three Fourier plots with x-axes spanning 0 to 40 Hz, and y-axes spanning 0 to 2 in magnitude.
  - Monophasic: The line monotonically decreases from 1.9 at 0 Hz to 0 at 40 Hz.
  - Biphasic (1:1): The line increases from 0.4 at 0 Hz to a peak of 1.8 at 5 Hz, and then decreases to 0 at 40 Hz.
  - Biphasic (2:1): The line monotonically decreases from 1.5 at 0 Hz to 0 at 40 Hz.
C. Three line plots with simulated responses to an alternating dark and light flash, with x-axes spanning 0 to 0.5s, and y-axes spanning −4 to 4 in response amplitude. The dark flash lasts from 0 to 0.25s, and the light flash lasts from 0.25 to 0.5s.
  - Monophasic: The line increases from 0 to 2 between 0-0.05s, stays at 2 until 0.25s, and decreases back to 0 between 0.25-0.3s.
  - Biphasic (1:1): The line increases from 0 to 2 between 0-0.05s, and decreases gradually back to 0 between 0.05-0.25s. It then decreases from 0 to −2 between 0.25-0.3s, and increases gradually back to −0.2 between 0.3-0.5s.
  - Biphasic (2:1): The line increases from 0 to 2 between 0-0.05s, and decreases sharply past 0 to −2 between 0.05-0.25s. It then decreases from −2 to −4 between 0.25-0.3s, and increases to −0.5 between 0.3-0.5s.

<u>Back to Fig. S2</u>

**Figure S3 Long Description**

A. Two panels of line plots showing modeled L1 responses to a moving light or dark edge. Each panel has six lines showing modeled L1 responses to motion blurred edges with varying *g* values, (*g* = 0, 0.5, 1), modeled L1 responses without feedback, and the input stimuli. The y-axis is Activation (A.U.) spanning −1 to 1, and the x-axis is time (s) spanning 0 to 0.8 second. Values provided here are approximate.
  - Moving light edge
    ∘ The input is a dotted line that starts at 0, increases to 1 at 0.15s, and then decreases to −0.5 at 0.62s. (This is sign-inverted from the examples in Fig. 3C.)
    ∘ The blurred input is a dotted line that matches the input, except the two changes (from 0 to 1, and from 1 to −0.5) are slower and smoother, taking 0.1s to transition rather than being instantaneous.
    ∘ The model-without-feedback trace and the g = 0 trace are both shaped similarly to the blurred input trace, but are shifted later. Like the input, they also travel from 0 to 1 to −0.5 A.U. The transition from 0 to 1 occurs at 0.18-028s, and the transition from 1 to −0.5 occurs at 0.65-0.75s. The g = 0 trace is slightly different at the end of the stimulus, where it stops decreasing at −0.25 and begins to return to 0, whereas the model-without feedback trace continues to decrease until it reaches −0.5 like the stimulus does.
    ∘ The g = 1 trace differs in that it initially overlaps with the g = 0 trace, but instead of going all the way up to 1, it stops at 0.25 and goes back down, reaching 0.1 at around 0.3s. It stays at 0.1 until 0.65s, and then decreases to a trough of −1 at 0.75s.
    ∘ The g = 0.5 trace is intermediate between the g = 0 and g = 1 traces (but is closer to the g = 1 trace).
  - Moving dark edge
    ∘ The input is a dotted line that starts at 0, drops to −1 at 0.15s, and then jumps to 0.5 at 0.62s. (This is the same as Fig. 3C.)
    ∘ The blurred input is a dotted line that matches the input, except the two changes (from 0 to −1, and from −1 to 0.5) are slower and smoother, taking 0.1s to transition rather than being instantaneous.
    ∘ The model-without-feedback trace and the g = 1 trace are both shaped similarly to the blurred input trace, but are shifted later. Like the input, they also travel from 0 to 1 to −0.5 A.U. The transition from 0 to 1 occurs at 0.18-028s, and the transition from 1 to −0.5 occurs at 0.65-0.75s. The g = 1 trace is slightly different at the end of the stimulus, where it stops decreasing at −0.25 and begins to return to 0, whereas the model-without feedback trace continues to decrease until it reaches −0.5 like the stimulus does.
    ∘ The g = 0 trace differs in that it initially overlaps with the g = 1 trace, but instead of going all the way down to 1, it stops at 0.25 and goes back up, reaching −0.1 at around 0.3s. It stays at −0.1 until 0.65s, and then increases to a peak of 1 at 0.75s.
    ∘ The g = 0.5 trace is intermediate between the g = 1 and g = 0 traces (but is closer to the g = 0 trace).
  - An arrow above both plots at around 0.65 emphasizes the difference between the traces at that time point, where the impact of the variably engaged feedback levels is apparent. By that time point, the light edge has engaged feedback when g = 1 (or 0.5), altering their responses to stimuli at 0.65s relative to the model without feedback. In contrast, the dark edge responses show evidence of engaged feedback when g = 0 (or 0.5).
B. Same as A, but for L2. The input, blurred input, and model-without-feedback traces are the same as in A. The model-without-feedback trace still overlaps with the g = 0 trace for the light edge, and the g = 1 trace for the dark edge. The g = 0.5 trace is also still intermediate between the *g* = 0 and *g* = 1 traces. An arrow is also above both plots at around 0.65 to emphasize the difference between the traces at that time point, where the impact of the variably engaged feedback levels is apparent.
  ∘ Moving light edge: The g = 1 trace stops increasing a bit higher than in A, at 0.5 (instead of 0.25). After the second transition, it decreases to a larger trough of −1.2 (instead of −1).
  ∘ Moving dark edge: The g = 0 trace stops decreasing a bit lower than in A, at −5 (instead of −0.25). After the second transition, it increases to a larger peak of 1.2 (instead of 1).

<u>Back to Fig. S3</u>

**Figure S4 Long Description**

The x-axis of these plots is time (s) spanning 0 to 0.8s. The y-axis of these plots is Activation (A.U) spanning −1 to +1.

A. Input: Line plot showing the same moving light edge and moving dark edge stimuli as in Fig. S3 overlaid together. The light edge starts at 0, increases to +1, and then decreases to −0.5. The dark edge starts at 0, decreases to −1, and then increases to +0.5.
B. L1 Pathway: Line plot showing modeled L1 responses to the stimuli in A. Responses greater than zero are shown as more saturated/darker than responses less than zero to distinguish between responses that are known to be relevant to downstream neurons after half-wave rectification has occurred.
  - Dotted lines show responses from the model without feedback. The light response transitions from 0 to +1, remains at +1, and then transitions to −0.5. The dark response transitions from 0 to −1, remains at −1, and then transitions to +0.5. The saturated (>0) and unsaturated (<0) portions of this set of responses are symmetric.
  - Solid lines show responses from the full model with feedback. The light response transitions from 0 to +1, decreases slightly to +0.8, and then transitions to −0.25, following a similar time course as the dotted no-feedback light response. The dark response transitions from 0 to −0.4, turning around sooner than the no-feedback dark response and staying at −0.2; it then increases faster and more strongly to +0.8 compared to the no-feedback dark response. The saturated (>0) and unsaturated portions of this set of responses is not symmetric, and notably the feedback model of the saturated portion has a faster response at 0.65s than the no-feedback model.
C. L2 Pathway: Line plot showing modeled L2 responses to the stimuli in A. Responses greater than zero are shown as more saturated/darker than responses less than zero to distinguish between responses that are known to be relevant to downstream neurons after half-wave rectification has occurred.
  - Dotted lines show responses from the model without feedback, which are sign inverted relative to the L1 pathway in B. The light response transitions from 0 to −1, remains at −1, and then transitions to +0.5. The dark response transitions from 0 to +1, remains at +1, and then transitions to −0.5. The saturated (>0) and unsaturated (<0) portions of this set of responses are symmetric.
  - Solid lines show responses from the full model with feedback. The light response transitions from 0 to −0.75, increases gradually to −0.4, and then transitions to +0.5, following a faster time course than the no-feedback light response. The dark response transitions from 0 to +0.5, turning around sooner than the no-feedback dark response and staying at +0.1; it then increases faster and more strongly to −1 compared to the no-feedback dark response. The saturated (>0) and unsaturated portions of this set of responses is not symmetric, though the feedback model of both portions have faster responses at 0.65s than the no-feedback model.

<u>Back to Fig. S4</u>

**Figure S5 Long Description**

Two panels labeled High speed and Low speed, each with three violin plots displaying the distribution of temporal offsets generated from the model without feedback, the full feedback model, and the model with increased feedback. The x-axes are of temporal offset in milliseconds, ranging from −160 to 160. The following describes the violin plots with statistics rounded to the nearest digit:

- High speed:
  ∘ The model without feedback has quartiles at −92, −80, and −71, and a range from −156 to −55.
  ∘ The full feedback has quartiles at −53, −45, and −28, and a range from −149 to 156.
  ∘ The model with increased speed has quartiles at −43, −34, and −16, and a range from −131 to 156.
  ∘ All three distributions are significantly different from each other (***, p<0.001).
- Low speed:
  ∘ The model without feedback has quartiles at −47, −45, and −44, and a range from −57 to −38.
  ∘ The full feedback has quartiles at −32, −25, and −3, and a range from −68 to 156.
  ∘ The model with increased speed has quartiles at −27, −16, and 17, and a range from −50 to 156.
  ∘ All three distributions are significantly different from each other (***, p<0.001).

<u>Back to Fig. S5</u>

**Figure S6 Long Description**

A. Series of three images. The first is a panoramic color photo of forest scenery, consisting mostly of tree trunks at various distances. The second is the same photo after removing gamma correction and converting to grayscale. The third is a further modified version of the photo after applying optical blurring.
B. Line plot in which the y-axis is yaw velocity (degrees per second) ranging from −400 to 400, and the x-axis is time (seconds) ranging from 0 to 2. A single trace fluctuates smoothly throughout most of the range of yaw velocities over time, beginning at approximately −50 degrees/second at time 0. The trace has four prominent peaks at approximately (−250, 0.2), (400, 0.6), (−200, 1.1), (300, 1.7), with several smaller local extrema in between the four peaks.
C. Line plot with three overlaid traces and scale bars for the x- and y-axes. The y-axis is in arbitrary units, spanning approximately 0.08 a.u., and the x-axis is time, spanning 2 seconds. The first trace is an inverted, normalized version of the naturalistic stimulus (see Fig. 6A for description). The second trace is a normalized version of the measured response of L2. The third trace is the output of the full recurrent feedback model after fitting it to the naturalistic stimulus. The measured L2 response and the model output are closely correlated (r^2^ = 0.90), and the two traces both generally follow the trajectory of the stimulus, lagging slightly behind in time.
D. Same as C, except the third trace is the output of the model without feedback. This trace is less closely correlated with the measured L2 response (r^2^ = 0.56) than in C, and at times it lags noticeably behind not only the stimulus but also the measured L2 response.

<u>Back to Fig. S6</u>

